# Single-cell generalized trend model (scGTM): a flexible and interpretable model of gene expression trend along cell pseudotime

**DOI:** 10.1101/2021.11.25.470059

**Authors:** Elvis Han Cui, Dongyuan Song, Weng Kee Wong, Jingyi Jessica Li

## Abstract

**Motivation:** Modeling single-cell gene expression trends along cell pseudotime is a crucial analysis for exploring biological processes. Most existing methods rely on nonparametric regression models for their flexibility; however, nonparametric models often provide trends too complex to interpret. Other existing methods use interpretable but restrictive models. Since model interpretability and flexibility are both indispensable for understanding biological processes, the single-cell field needs a model that improves the interpretability and largely maintains the flexibility of nonparametric regression models.

**Results:** Here we propose the single-cell generalized trend model (scGTM) for capturing a gene’s expression trend, which may be monotone, hill-shaped, or valley-shaped, along cell pseudotime. The scGTM has three advantages: (1) it can capture non-monotonic trends that are still easy to interpret, (2) its parameters are biologically interpretable and trend informative, and (3) it can flexibly accommodate common distributions for modeling gene expression counts. To tackle the complex optimization problems, we use the particle swarm optimization algorithm to find the constrained maximum likelihood estimates for the scGTM parameters. As an application, we analyze several single-cell gene expression data sets using the scGTM and show that it can capture interpretable gene expression trends along cell pseudotime and reveal molecular insights underlying the biological processes.

**Availability and implementation:** The Python package scGTM is open-access and available at https://github.com/ElvisCuiHan/scGTM.

## 1 Introduction

Pseudotime analysis is one of the most important topics in single-cell transcriptomics. There has been fruitful work on inferring cell pseudotime (Magwene et al., 2003; Bendall et al., 2014; Trapnell et al., 2014; Shin et al., 2015; Ji and Ji, 2016; Qiu et al., 2017; Street et al., 2018; Mondal et al., 2021; Cao et al., 2019) and constructing statistical models for gene expression along the inferred cell pseudotime (Campbell and Yau, 2017; Bacher et al., 2018; Van den Berge et al., 2020; Ren and Kuan, 2020; Song and Li, 2021). Informative trends of gene expression along cell pseudotime may reflect molecular signatures in the biological processes. For instance, a gene may over time exhibit a *hill-shaped* trend (i.e. first-upward-then-downward) (Fig. 1b) or a *valley-shaped* trend (i.e. first-downward-then-upward) (Fig. 1c) trend and either of the trend may indicate the occurrence of some biological event. Hence, it is of great interest to have a statistical model that can capture the different gene expression trends accurately along cell pseudotime.

**Fig. 1.**
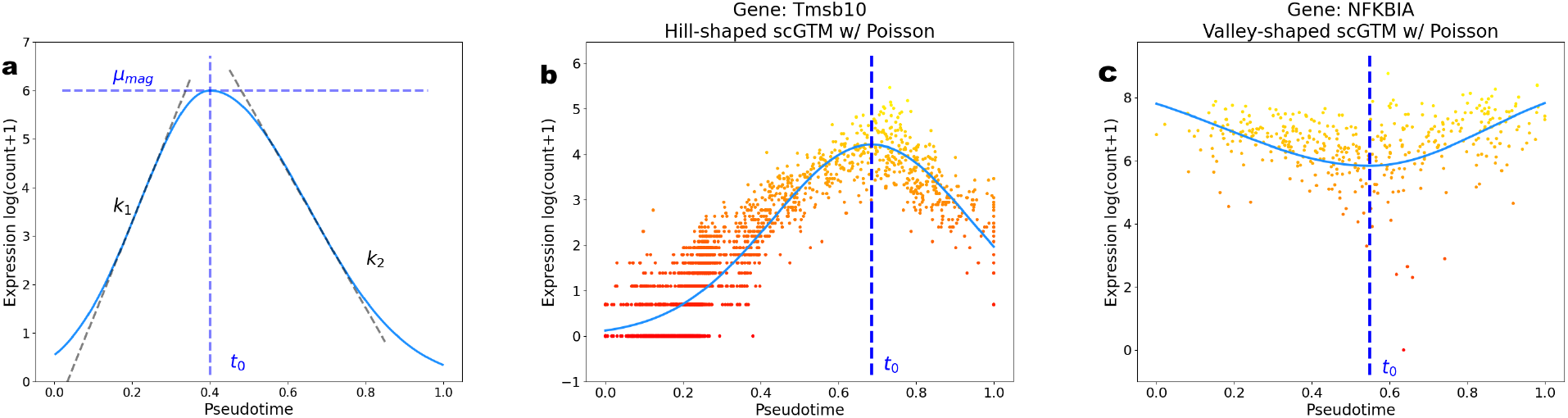
Illustration of the scGTM. (**a**) Four parameters of the scGTM in Equation (2) for a hill-shaped trend: the maximum log expected expression *μ*_mag_ (horizontal blue line), the activation strength *k*_1_ (absolute value of the left tangent line’s slope), the repression strength *k*_2_ (absolute value of the right tangent line’s slope), and the change time *t*_0_ (vertical blue line). (**b**) A hill-shaped trend of gene *Tmsb10* (in the GYRUS dataset) fitted by the scGTM with counts modeled by the Poisson distribution. (**c**) A valley-shaped trend of gene *NFKBIA* (in the LPS dataset) fitted by the scGTM with counts modeled by the Poisson distribution. In b–c, the scatter points indicate gene expression levels, and the curves are the trends fit by the scGTM.

There are two types of statistical methods that have been developed to model the relationship between a gene’s expression in a cell (or a sample) and the cell pseudotime. The latter is the sample’s physical time and so modeling issues encountered here are similar to those commonly seen in fitting a regression model to a data set in statistics. Methods of the first type are based on statistical regression models, such as generalized linear models (GLM) and generalized additive models (GAM), where frequently parameters in these models do not have direct relevance to gene expression dynamics. Specifically, the GLM used in the Monocle3 method (Cao et al., 2019) assumes that a gene’s log-transformed expected expression in a cell is a linear function of the cell pseudotime, making it unable to capture hill- and valley-shaped trends that linear trends cannot approximate well. Consequently, most methods use nonparametric regression models, such as the GAM and piece-wise linear models, to capture complex trends. For example, Storey et al. (2005) applied basis regression; Trapnell et al. (2014) considered the GAM with the Tobit likelihood; Ren and Kuan (2020) applied the GAM with Bayesian shrinkage dispersion estimates; Van den Berge et al. (2020) proposed tradeSeq using the spline-based GAM. More recently, Song and Li (2021) proposed the PseudotimeDE method, which fixes the p-value calibration issue in tradeSeq and also uses the spline-based GAM with spline functions. Additionally, Bacher et al. (2018) used a piecewise linear model, which is more restrictive than the GAM. Although these nonparametric regression methods can fit complex gene expression trends, they are prone to over-fitting without proper hyperparameter tuning (as we will show in Section 3) and their parameters either do not directly inform the shape of a trend (e.g., hill-shaped) or carry biological meanings.

Unlike the first type, methods of the second type use models with direct relevance to gene expression dynamics, and notable methods include ImpulseDE/ImpulseDE2 (Chechik and Koller, 2009; Sander et al., 2017; Fischer et al., 2018) and switchDE (Campbell and Yau, 2017). Specifically, ImpulseDE2 estimates a gene expression trend using a double-logistic curve to capture the non-monotone trends; however, even though the parameters have biological interpretations, they do not intuitively inform the shape of a trend. In contrast, switchDE uses a restrictive model with parameters that directly inform the shape of a trend (e.g., a gene’s activation time) but is unable to detect non-monotonic trends.

The above review suggests that there is no current model that can capture monotone, hill-shaped, and valley-shaped trends with biologically interpretable and trend-informative parameters. To this end, we propose the scGTM that (i) can capture both hill- and valley-shaped trends and monotone trends, (ii) has interpretable and trend-informative parameters, and (iii) has flexible modeling for count data.

To estimate the scGTM parameters, we apply particle swarm optimization (PSO) to find the constrained maximum likelihood estimates (MLE) of the model parameters (Fig. S21). PSO has several advantages that make it suitable for our optimization problem: (i) it does not require the objective function to be convex or differentiable; (ii) it can handle boundary constraints and discrete parameters without having to re-formulate the objective function, and (iii) unlike the Newton-type algorithms used in Trapnell et al. (2014); Wood (2017); Campbell and Yau (2017), PSO is gradient-free. In addition, PSO codes are widely and freely available, easy to implement and its successes in tackling complex optimization problems are already well documented in computer science and engineering.

The rest of the paper is organized as follows. In Section 2, we introduce the scGTM and briefly review the PSO algorithm. In Section3, we compare the scGTM with the GLM, GAM, ImpulseDE2, and switchDE and show its advantages in capturing informative, interpretable gene expression trends in two real data sets. Section 4 contains a discussion and future work.

## 2 Methods

### 2.1 The scGTM formulation

Let **Y** = (*y_gc_*) be an observed *G* × *C* gene expression count matrix, where *G* is the number of genes, *C* is the number of cells (i.e., the number of pseudotime values), and *y_gc_* is the (*g*, *c*)-th element indicating the observed expression count of gene *g* = 1,…, *G* in cell *c* = 1,…, *C*. We consider gene expression counts as random variables whose randomness comes from experimental measurement uncertainty, so *y_gc_* is a realization of the random count variable *Y_gc_*. Given a particular gene *g*, for notation simplicity, we drop the subscript *g* and denote *Y_gc_* as *Y_c_* and *y_gc_* as *y_c_*. We denote by *t_c_* ∈ [0, 1] the inferred (normalized) pseudotime of cell *c*. In the scGTM, *t*_1_,…,*t_C_* are treated as fixed values of pseudotime and serve as the covariate vector of interest.

Given *t_c_*, the scGTM can model the count variable *Y_c_* using four count distributions commonly used for gene expression data: the Poisson, negative binomial (NB), zero-inflated Poisson (ZIP), and zero-inflated negative binomial (ZINB) distributions.

For a hill-shaped gene, the scGTM is

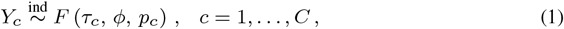

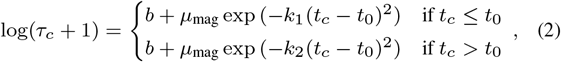

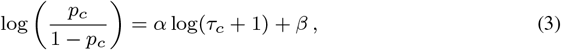

where *F* (*τ_c_*, *ϕ*, *p_c_*) in (1) represents one of the four common count distributions. The most general case is when *F* (*τ_c_*, *ϕ*, *p_c_*) = ZINB (*τ_c_*, *ϕ*, *p_c_*) with mean parameter *τ_c_* ≥ 0, dispersion parameter 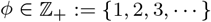 and zero-inflated parameter *p_c_* ∈ [0, 1].As special cases, *F* (*τ_c_*, *ϕ*, 0) = NB (*τ_c_*, *ϕ*), *F* (*τ_c_*, ∞, *p_c_*) = ZIP (*τ_c_*, *p_c_*), and *F* (*τ_c_*, ∞, 0) = Poisson (*τ_c_*).

We design the parametric form (2) for the following reasons. First, on the left-hand side, log(*τ_c_* + 1) is motivated by the logarithmic link function used in the generalized linear models (GLM) and GAM. The addition of 1 is to ensure that log(*τ_c_* + 1) ≥ 0 so we can use *b* ≥ 0 as the baseline of the hill-shaped trend (empirically, b is set to 0 and works well). Second, on the right-hand side, two partial Gaussian functions are adopted to model the trend’s increasing and decreasing parts separately, so that the trend is allowed to be asymmetric (e.g., the increasing trend may be steeper or flatter than the decreasing trend). We choose to piece two partial Gaussian functions at the maximum into one function for two reasons: (1) the function is smooth (differentiable everywhere) and has a zero derivative at the maximum; (2) the function has tails that converge to the baseline b as the pseudotime *t*_c_ moves away from the mode *t*_0_, a pattern that agrees with many biological processes.

Fig. 1a shows the roles of the four parameters *μ*_mag_, *k*_1_, *k*_2_, and *t*_0_ in (2) for modelling a hill-shaped trend. For a valley-shaped trend, there are four similar parameters and we note that a monotone increasing trend is a special case of a hill-shaped trend with the increasing part only. The four parameters in the Fig. 1a are the maximum log expected expression *μ*_mag_, the activation strength *k*_1_, the repression strength *k*_2_, and the change time *t*_0_ where the expected expression stops increasing. Fig. 1b–c show the scGTM fits to the gene (*Tmsb10* in the GYRUS data set and another gene *NFKBIA* in the LPS data set from the Supplementary Table S1). Their trends reveal a hill- and valley-shaped trend., respectively.

In this hill-shaped scGTM, we assume that the gene’s expression count *Y_c_* in cell *c* has mean parameter *τ_c_* and zero-inflation parameter *p_c_*, and both depend on the pseudotime *t_c_* of cell *c*. In (2), we link *τ_c_* to *t_c_* by assuming that log(*τ_c_* + 1) is a non-negative transformation that compresses extremely large values of *τ_c_* using a two-part Gaussian function corresponding to *t_c_* ≤ *t*_0_ and *t_c_* > *t*_0_; we choose the Gaussian function for its good mathematical properties and interpretability. We link *p_c_* to *t_c_* in (3) using a logistic regression, with predictor log(*τ_c_* + 1), i.e., the logistic transformation of *p_c_* is a linear function of log(*τ_c_* + 1) (with slope *α* and intercept *β*) and thus a function of *t_c_*.

Besides 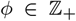 and *α*, 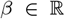, the following parameters of the hill-shaped scGTM shown in Fig. 1a need to be estimated for biological interpretations:

- *μmag* ≥ 0: magnitude of the hill, i.e., *μmag* = max_*c*∈{1,…,*C*}_ log(*τ_c_* + 1);
- *k*_1_ ≥ 0: activation strength (how fast the gene is up-regulated);
- *k*_2_ ≥ 0: repression strength (how fast the gene is down-regulated);
- *t*_0_ ∈ [0, 1]: change time (where the gene reaches the maximum expected expression). It is within [0, 1] since the pseudotime has been re-scaled.

For a valley-shaped gene, the scGTM is the same except that we replace (2) by

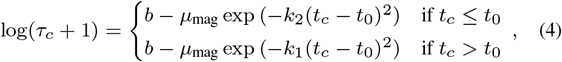

where *b* indicates the baseline (maximum) log-transformed (expected expression +1) of the valley-shaped gene. The interpretation of the four key parameters of the valley-shaped scGTM becomes

- *μ*_mag_ ∈ [0, *b*]: magnitude of the valley, i.e., *b* – *μ*_mag_ = min_*c*∈{ι,…,*C*}_ log(*τ_c_* + 1);
- *k*_1_ ≥ 0: activation strength (how fast the gene is up-regulated);
- *k*_2_ ≥ 0: repression strength (how fast the gene is down-regulated);
- *t*_0_ ∈ [0,1]: change time (where the gene reaches the minimum expected expression).

Compared to the hill-shaped scGTM, the valley-shaped scGTM has an additional baseline parameter b that needs to be estimated. For simplicity, we estimate *b* by a plug-in estimator 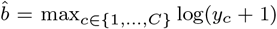, where *y*_1_,…,*y_C_* are the observed counts of a valley-shaped gene. This estimate is justified by the fact that the maximum likelihood estimate (MLE) of the upper bound parameter of a domain is the maximum order statistic; i.e., if *x*_1_,…,*x_n_* are randomly sampled from a distribution with domain [*a*, *b*], then 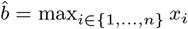 is the MLE of b. For the common parameters of the hill- and valley-shaped scGTMs in Section 2.2, we next discuss how PSO can provide constrained likelihood estimates for these parameters.

### 2.2 Constrained MLE and the PSO algorithm

To fit the scGTM to a gene, we first need to ascertain whether the gene is hill- or valley-shaped: we first need to ascertain whether the gene is hill- or valley-shaped: we fit both hill- and valley-shaped models to the gene’s data and choose the model that has the smaller Akaike information criterion (AIC) value (see Supplementary Information S9 for the model selection results for the two genes in Fig. 1b–c). Next, based on the trend shape, we estimate the scGTM parameters. For a hill-shaped gene, we estimate the scGTM parameters Θ = (*μ*_mag_, *k*_1_, *k*_2_, *t*_0_, *ϕ*, *α*, *β*)^T^ from the observed expression counts ***y*** = (*y*_1_,…,*y_C_*)^T^ and cell pseudotimes ***t*** = (*t*_1_,…,*t_C_*)^T^ using the constrained maximum likelihood method, which respects each parameter’s range and ensures the estimation stability. Let log *L*(Θ | ***y**, **t***) be the log likelihood function and the optimization problem is:

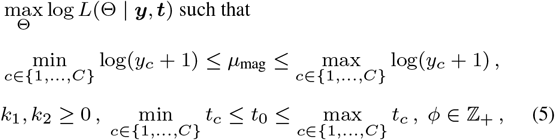

where

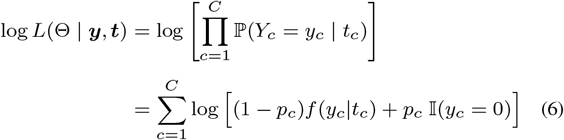

and

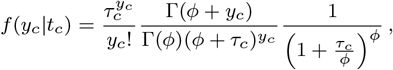

which can be further specified as a function of Θ based on (2) and (3).

For a valley-shaped gene, the constrained MLE problem is similar, and we omit the discussion for space consideration.

There are two difficulties in the optimization problem (5). First, the likelihood function (6) is neither convex nor concave. Second, the constraint is linear in *μ*_mag_, *k*_1_, *k*_2_, and *t*_0_ but *ϕ* is a positive integer-valued variable. Hence, conventional optimization algorithms such as P-IRLS in GAM (Wood, 2011, 2017) and L-BFGS in switchDE (Van Loan and Golub, 1996; Campbell and Yau, 2017) are difficult to apply in this case. Metaheuristics is a class of assumptions-free general purpose optimization algorithms that is widely and increasingly used to tackle challenging and high-dimensional optimization problems in the quantitative sciences (Whitacre, 2011a,b; Yang, 2017). PSO is an exemplary metaheuristic algorithm, and it has effectively solved various types of optimization problems. Korani and Mouhoub (2021) is a recent review of metaheuristic algorithms and their applications across various disciplines.

PSO first generates a swarm of candidate solutions (known as particles) to the optimization problem (5). At each iteration, particles change their positions within the constraints, and the algorithm finds the best solution among all particle trajectories. We summarize the vanilla PSO algorithm (Bratton and Kennedy, 2007) for the constrained MLE of the scGTM in Algorithm 1, and provide further details of PSO in the Supplementary Information.

#### Algorithm 1 PSO for the constrained MLE for the scGTM

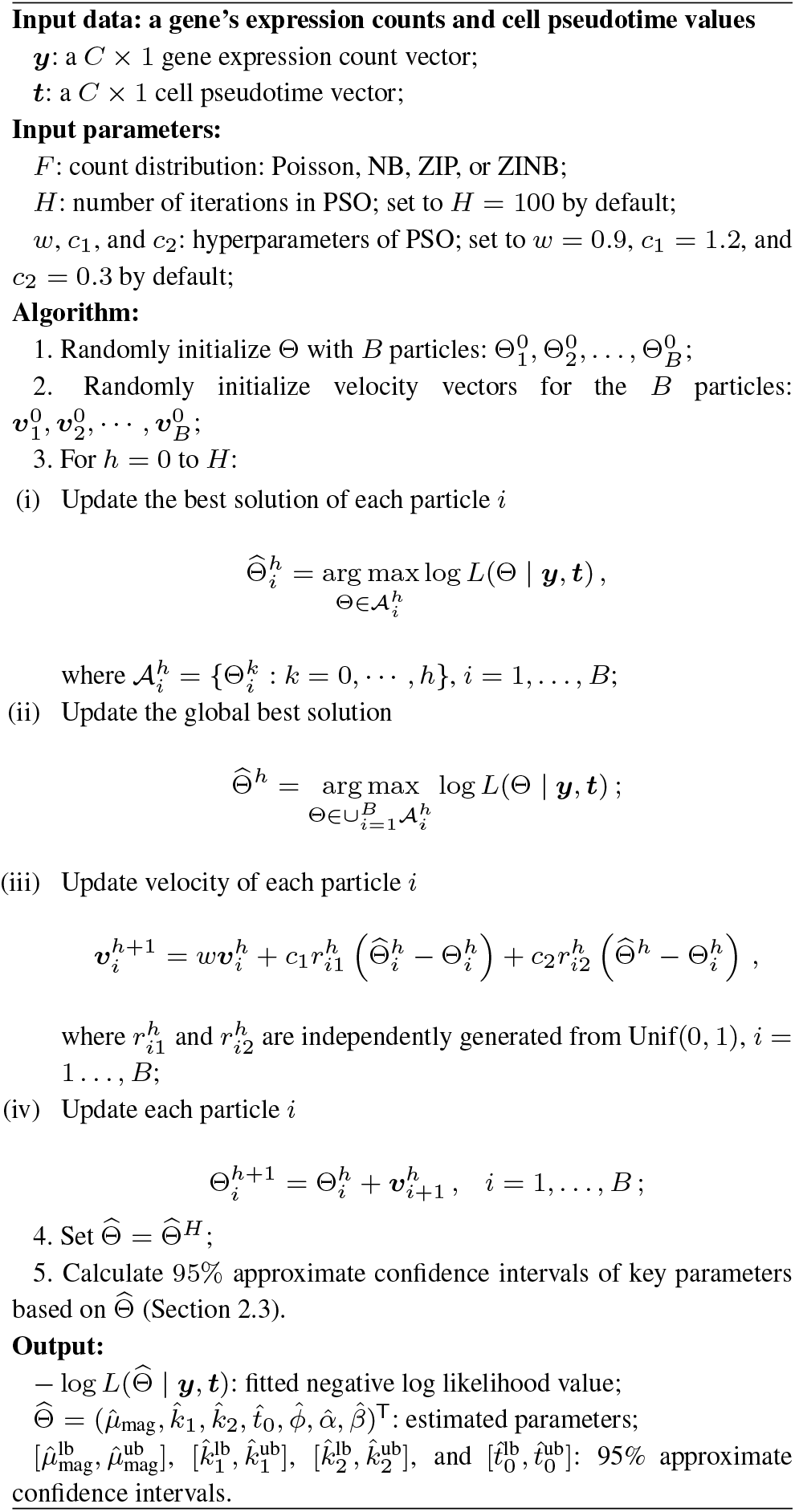

### 2.3 Approximate confidence intervals of the four key parameters in the scGTM

The estimated parameters 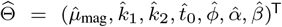 are next used to construct approximate confidence intervals for *μ*_mag_, *k*_1_, *k*_2_, and *t*_0_ using the maximum likelihood theory. Specifically, we calculate the plug-in asymptotic covariance matrix of 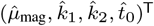 as the inverse of the partial Fisher information matrix of the four parameters evaluated at 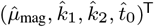 (detailed derivation in the Supplementary Information). Then we use the diagonal elements of this matrix as the plug-in asymptotic variances of 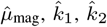, and 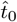, and denote them by 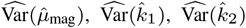, and 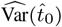, respectively. We then obtain a 95% approximate confidence interval for each of the parameters: 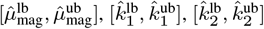, and 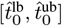, where

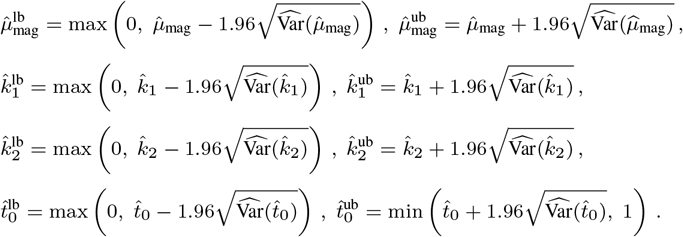

## 3 Results

### 3.1 scGTM outperforms GAM, GLM, LOESS, switchDE, and ImpulseDE2 in capturing informative and interpretable trends

As an example, we use the *MAOA* gene in the WANG dataset (Wang et al., 2020) (Supplementary Table S1) to compare the fitted trends of the scGTM, GAM, GLM, LOESS, switchDE, and ImpulseDE2. In the original study, the gene was reported to have a hill-shaped trend. Our comparison results have several interesting observations. First, we show that the scGTM provides more informative and interpretable gene expression trends than the GAM and GLM do when the count outcome comes from the Poisson, ZIP, NB, and ZINB distributions. Fig. 2a shows that the scGTM robustly captures the hill-shaped trends for the four distributions and consistently estimates the change time around 0.75, which is where the MAOA gene reaches its expected maximum expression. While the GAM also estimates the maximum expression around 0.75, its estimated trends are much more complex. This is likely due to possible overfitting (despite the use of penalization) and consequently, more difficult to interpret them than the scGTM trends (Fig. 2b). Unlike the scGTM and GAM, the GLM only allows for capturing monotone trends, making it unable to detect the possible existence of expression change time (Fig. 2c). Second, we compare the scGTM with the two existing methods, switchDE and ImpulseDE2, that use models with direct relevance to gene expression dynamics. Although switchDE estimates the activation time around 0.75, similar to the scGTM’s estimate change time, it cannot capture the downward expression trend as the cell pseudotime approaches 1.00 due to its monotone nature (Fig. 2d). ImpulseDE2 can theoretically capture a hill-shaped trend, but it only fits a monotone increasing trend for the *MAOA* gene (Fig. 2e). A likely reason is that the method was designed for time-course bulk RNA-seq data. Third, we compare the scGTM with the LOESS method commonly used for exploratory data analysis. While LOESS outputs a reasonable, though less smooth trend (Fig. 2f), it is not probability-based and thus does not have a likelihood. Hence, LOESS does not allow likelihood-based model selection, a functionality of the scGTM. To summarize, the scGTM fits outperform those from GAM, GLM, LOESS, switchDE and ImpulseDE2 by providing a more informative and interpretable trend with less concern on model overfitting.

**Fig. 2.**
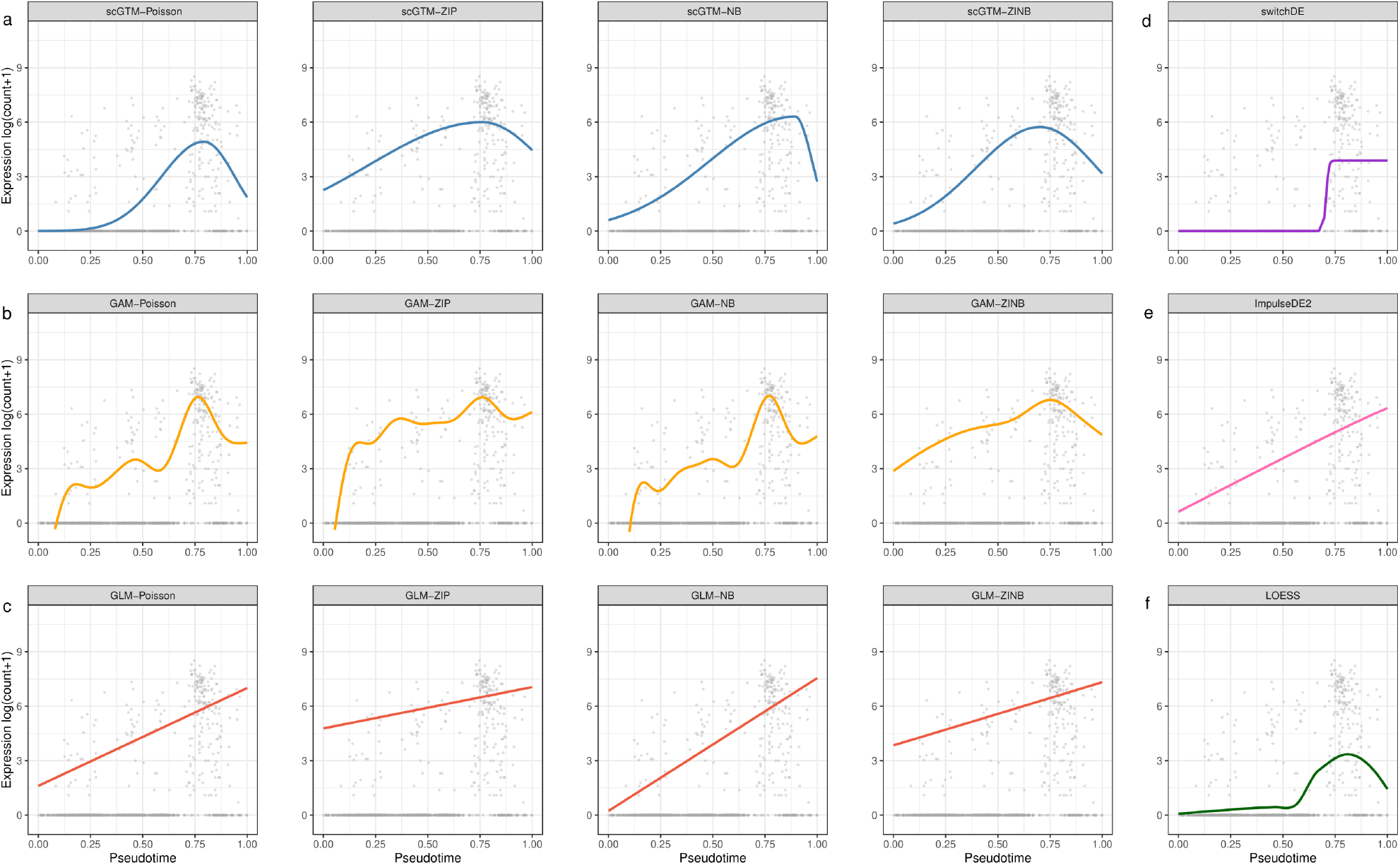
Comparison of the scGTM with GAM, GLM, LOESS, switchDE, and ImpulseDE2 for fitting the expression trend of gene *MAOA* in the WANG dataset (Wang et al., 2020) (Supplementary Table S1). In the first four columns, the three rows correspond to (a) scGTM, (b) GAM, and (c) GLM. From left to right, the first four columns correspond to Poisson, ZIP, NB, and ZINB as the count distribution used in the scGTM, GAM, and GLM. The fifth column corresponds to (d) switchDE, (e) ImpulseDE2 and (f) LOESS. Each panel shows the same scatterplot of gene *MAOA*’s log-transformed expression counts vs. cell pseudotime values, as well as a model’s fitted trend. With all four count distributions, the scGTM robustly captures the gene expression trend and estimates the change time around 0.75. In contrast, GLM, switchDE and ImpulseDE2 only describe the trend as increasing; GAM overfits the data and does not output trends as interpretable as the scGTM does; LOESS outputs a reasonable trend, but it does not allow likelihood-based model selection like the scGTM.

In addition to the *MAOA* gene, Wang et al. (2020) reported 19 other exemplary genes that define menstrual cycle phases and exhibit hill-shaped expression trends along the cell pseudotime. Supplementary Figs. S1-S19 compare the various model fits for the 19 genes and and we observe that the scGTM consistently provides more informative, interpretable trends than the other models.

Besides visually inspecting the fitted expression trends, we compare the AIC values of the scGTM, GAM, and GLM used with the four count distributions fitted to the aforementioned 20 genes. Note that a lower AIC value indicates a model’s better goodness-of-fit with the model complexity penalized. Supplementary Fig. S20 shows that the scGTM has comparable or even lower AIC values than the GAM’s AIC values, confirming that the scGTM fits well to data despite its much simpler model than GAM’s. Based on Fig. 2 and Supplementary Figs. S1–S20, we use the scGTM with the Poisson distribution in the following applications for its goodness-of-fit and model simplicity. This choice is consistent with previous research on modeling sequencing data (Silverman et al., 2020) and other count data (Warton, 2005; Campbell, 2021).

### 3.2 scGTM recapitulates gene expression trends of endometrial transformation in the human menstrual cycle

The WANG dataset contains 20 exemplar genes that exhibit temporal expression trends in unciliated epithelia cells in the human menstrual cycle Wang et al. (2020). The original study also ordered the 20 genes by the estimated pseudotime at which they achieved the maximum expression (Fig. 3a; genes ordered from top to bottom), and it was found that the ordering agreed well with the menstrual cycle phases (Fig. 3a; the top bar indicates the phases). Comparing the fitted expression trends of the 20 genes by the scGTM, switchDE, and ImpulseDE2, we observe that only the scGTM trends agree well with the data (Fig. 3). Additionally, we evaluate the 20 genes’ estimated change times (i.e., *t*_0_) by the scGTM and their estimated activation times by the switchDE. Although the change times and estimation times are both expected to correlate well with the gene ordering in the original study, only the change times estimated by the scGTM fulfills this expectation (Fig. 3b-c). Compared with the scGTM, switchDE miscalculates the activation times for many hill-shaped genes whose maximum expression occurs in the middle of the cycle; this is likely due to the fact that switchDE can only capture monotone trends (Fig. 3c). Similarly, ImpulseDE2 cannot well capture the trends of those hill-shaped genes (Fig. 3d). Unlike switchDE and ImpulseDE2, the scGTM estimates the change times reasonably for almost all genes. For instance, the *GPX3* gene has an estimated change time at 0.88, consistent with its role as a secretory middle/late phase marker gene Wang et al. (2020).

**Fig. 3.**
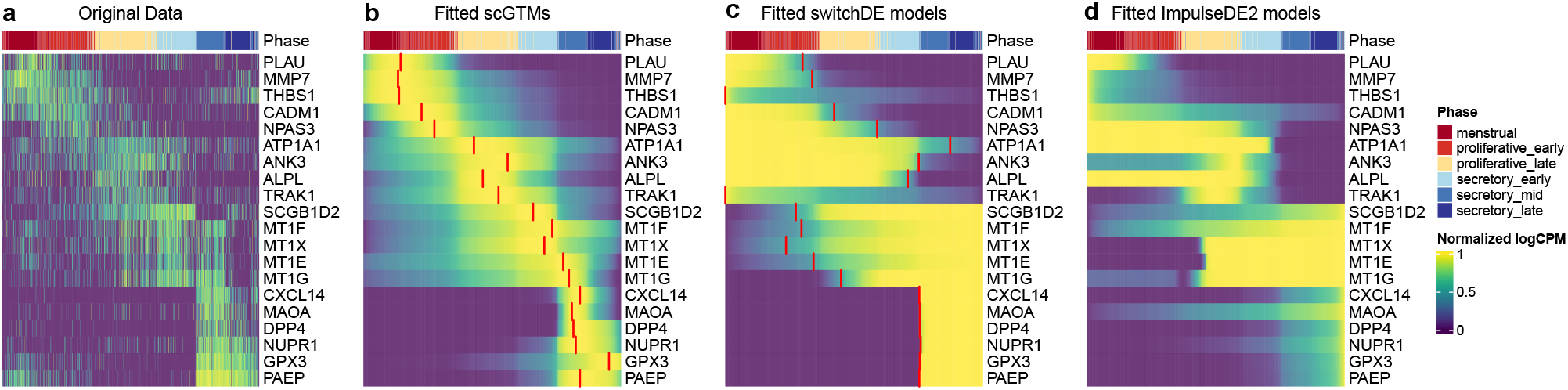
Fitted expression trends by the scGTM, switchDE, and ImpulseDE2 for 20 exemplar genes in the WANG dataset (Wang et al., 2020) (Supplementary Table S1). All panels are ordered by cell pseudotime values from 0 (left) to 1 (right). The top color bars show the endometrial phases defined in the original study. (a) The original expression values along pseudotime. (b) The fitted trends of the scGTM, with the red segments highlighting the estimated change times tθ. (c) The fitted trends of switchDE, with the red segments highlighting the estimated activation times. (d) The fitted trends of ImpulseDE2.

Besides the 20 exemplar genes, we apply the scGTM, switchDE, and ImpulseDE2 to fit the expression trends of all 1,382 menstrual cycle genes reported in Wang et al. (2020). Supplementary Fig. S28 shows that the scGTM still outperforms switchDE and ImpulseDE2 for capturing these genes’ expression trends. In summary, the scGTM provides useful summaries for gene expression trends in the human menstrual cycle.

### 3.3 scGTM identifies informative gene expression trends after immune cell stimulation

As the second real data application, we use the scGTM to categorize gene expression trends in mouse dendritic cells (DCs) after stimulation with lipopolysaccharide (LPS, a component of gram-negative bacteria) Shalek et al. (2014). First, we apply the likelihood ratio tests to screen the genes that the scGTM fits significantly better than the null Poisson model (in which *τ_c_* and *p_c_* in (1) do not depend on cell pseudotime *t_c_*). Assuming that the likelihood ratio statistic of every gene follows 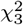 as the null distribution, we retain 2405 genes whose Benjamini-Hochberg (BH) adjusted *p*-values ≤ 0.01.

Second, we use the scGTM’s confidence levels of the three parameters *t*_0_, *k*_1_, and *k*_2_ to categorize the 2405 genes into three types: (1) *hill-shaped & mostly increasing genes*: 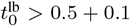 (change time occurs at late pseudotime) and 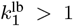 (strong activation strength), (2) *hill-shaped & mostly decreasing genes*: 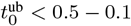 (change time occurs at early pseudotime) and 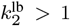 (strong repression strength), and (3) *valley-shaped genes*. To demonstrate that this categorization is biologically meaningful, we perform gene ontology (GO) analysis on the three gene types and compare the enriched GO terms. Fig. 4a shows that the three gene types are enriched with largely unique GO terms, verifying their functional differences. Notably, the hill-shaped & mostly increasing genes are related to immune response processes, showing consistency between their expression trends (activation after the LPS stimulation) and functions (immune response). Further, we visualize 5 illustrative genes from each gene type (Fig. 4b) and observe that the scGTM’s fitted trends agree well with the data. In conclusion, the scGTM can help users discern genes with specific trends by its trend-informative parameters.

**Fig. 4.**
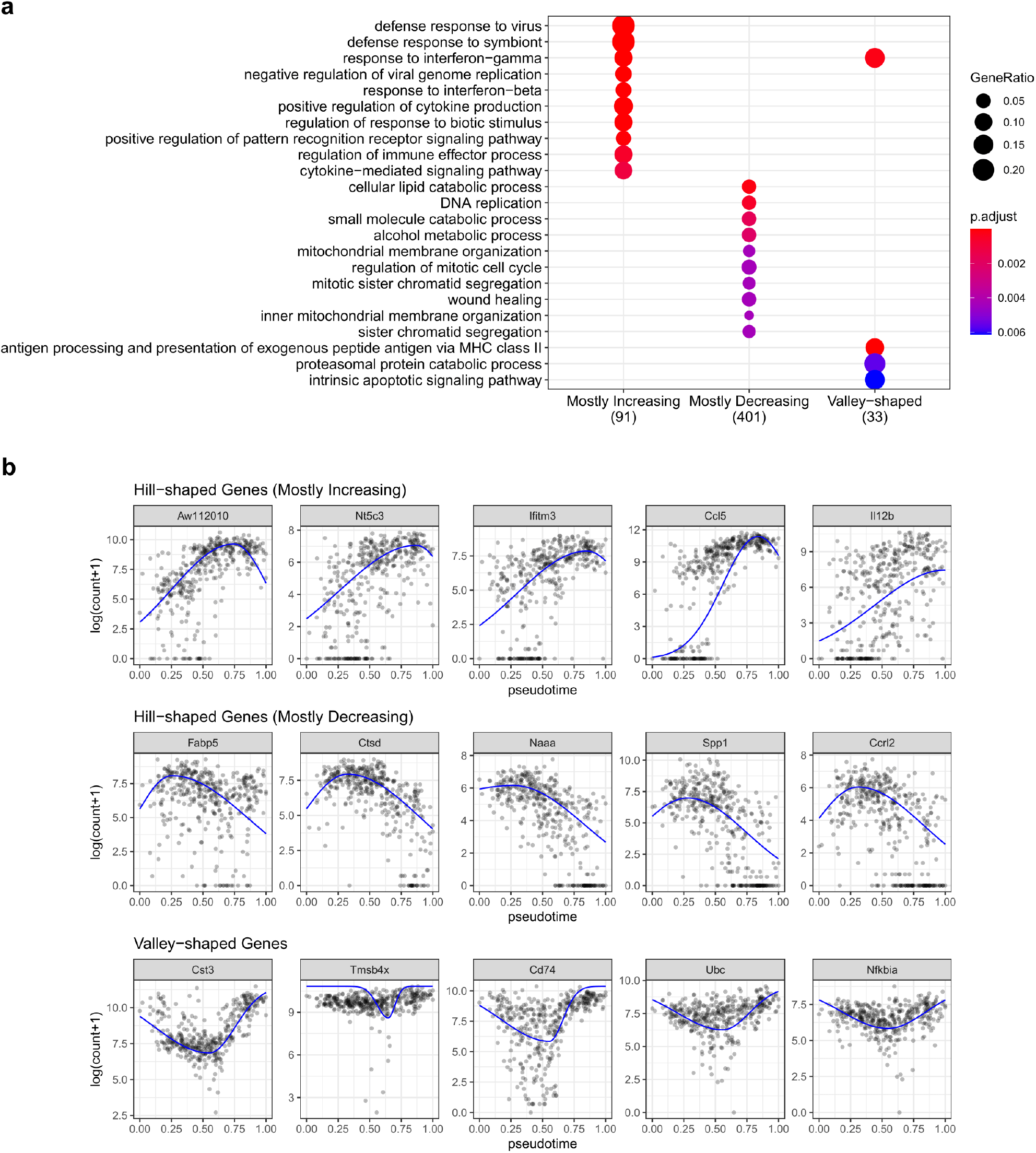
Three types of gene expression trends characterized by the scGTM parameters in the LPS dataset (Supplementary Table S1). (a) GO enrichment analysis of the three gene types. The top enriched GO terms are different among the three gene types. Notably, the hill-shaped & mostly increasing genes (1st column) are functionally related to immune responses. (b) Visualization of example genes in the three types. The scatter plots show gene expression data; the trends estimated by the scGTM (blue curves) well match the data.

Besides the above three real data applications, we conduct a simulation study to verify the robustness of the scGTM to gene expression trends not generated from the scGTM assumptions. The simulation results also show that, beyond good interpretability, the scGTM is flexible enough to fit various trends to a similar extent as the GAM does (Supplementary Information S3). Moreover, we use a bootstrap analysis to show that the fitted scGTM trend has a smaller variance than the fitted GAM trend does (Supplementary Information S3), at the cost of a larger bias.

## 4 Discussion

We propose the scGTM as a flexible and interpretable statistical model for studying single-cell gene expression trend along cell pseudotime. Using four count distributions and two real data sets, we demonstrated that the scGTM has interpretable parameters that can directly inform a trend for gene expression counts. The scGTM parameters are estimated by the constrained maximum likelihood estimation via PSO, which is one of the most popular metaheuristic algorithms for function optimization. We showed that scGTM has distinct advantages over the classic models GLM and GAM and the two recent methods switchDE and ImpulseDE2 in that it can uniquely capture robust, informative, and interpretable trends. In contrast, the GLM and switchDE can only estimate monotonic trends; the GAM often provides trends that are too complex to interpret, and ImpulseDE2 (a method designed for bulk RNA-seq data) does not have stable performance on single-cell data. The estimated parameters and confidence intervals from the scGTM are then used to characterize the expression trends of the genes.

Note that the scGTM is extendable by assuming a more complicated mean function, whose estimation can still be achieved by the PSO algorithm (whose major advantage is its flexibility). To demonstrate this functionality of the scGTM, we have added a simulation in Supplementary Information S10, where we use the sine function to generate one gene’s true expression trend along the pseudotime. With its mean function set as as the sine function, the scGTM accurately estimates the gene trend (Supplementary Fig. S25). In a future version of the scGTM package, we can allow users to input specified mean functions that reflect the gene expression trends they are interested in. On the other hand, if users do not have a prior preference for the gene expression trends, we would recommend them to use the generalized additive models that can capture flexible trends.

Strictly speaking, the inference of the scGTM has two caveats. First, the parameter estimation includes a double-dipping procedure: the same data are first used to decide whether a trend is hill- or valley-shaped and it is used to estimate the parameters. Second, since only the key parameters *μ*_mag_, *k*_1_, *k*_2_, and *t*_0_ are inferential targets, the other parameters *ϕ*, *α* and *β* should be regarded “nuisance” parameters. However, the construction of confidence intervals of the key parameters does not account for these two caveats and would thus result in overly optimistic confidence intervals. We will investigate how to obtain better-calibrated confidence intervals in future research.

In our previous work (Song and Li, 2021), we developed a method PseudotimeDE to account for the uncertainty of inferred pseudotime on the inference of differentially expressed genes along the pseudotime. Note that PseudotimeDE is directly extendable to the scGTM, by just replacing the GAM in PseudotimeDE by the scGTM. However, here our focus is on proposing the scGTM for interpreting a trend, instead of testing whether a trend is different from a horizontal line, i.e., the focus of PseudotimeDE. Hence, we leave the incorpration of the scGTM into PseudotimeDE to future work. Moreover, we have a simulation study to show that the fitted scGTM trends have shapes largely robust to noise added to pseudotime (**Supplementary Information S4**).

The current implementation of the scGTM is only applicable to a single pseudotime trajectory (i.e., cell lineage). A natural extension is to split a multiple-lineage cell trajectory into single lineages and fit the scGTM to each lineage separately.

In addition, the vanilla PSO algorithm in this paper handles each parameter’s constraint separately. Hence, if we need a constraint on more than one parameter, e.g., *k*_1_/*k*_2_ should be within a a user-specified range, then we have to develop a variant algorithm of PSO or use other metaheuristics algorithms.

## 5 Data Availability

The code and data for genearing the results are available at zenodo (doi: 10.5281/zenodo.5728342). The scGTM Python package is available at https://github.com/ElvisCuiHan/scGTM.

## 6 Acknowledgements

The authors would like to thank Dr. Wanxin Wang for providing the data in Wang et al. (2020). The authors also appreciate the comments and feedback from the members of the Junction of Statistics and Biology at UCLA (http://jsb.ucla.edu).

## 7 Funding

This work was supported by National Science Foundation DBI-1846216 and DMS-2113754, NIH/NIGMS R01GM120507 and R35GM140888, Johnson and Johnson WiSTEM2D Award, Sloan Research Fellowship, and UCLA David Geffen School of Medicine W.M. Keck Foundation Junior Faculty Award (to J.J.L.).

## Notes

https://github.com/ElvisCuiHan/scGTM

https://doi.org/10.5281/zenodo.5728341

### Competing Interest Statement

The authors have declared no competing interest.

### Summary of Updates

New simulation and real-data analysis results were added.

