## Supplementary for "Single-cell generalized trend model (scGTM): a flexible and interpretable model of gene expression trend along cell pseudotime"

#### Supplementary Information

#### S1 Fitted trends of 19 genes in the WANG dataset

In this section, we present the other 19 exemplar genes (in addition to *MAOA*) in the WANG dataset [Wang et al. (2020)] and their fitted trends by the scGTM, GAM, GLM, LOESS, switchDE, and ImpulseDE2. The interpretation of each figure is the same as Fig. 2 in the main text.

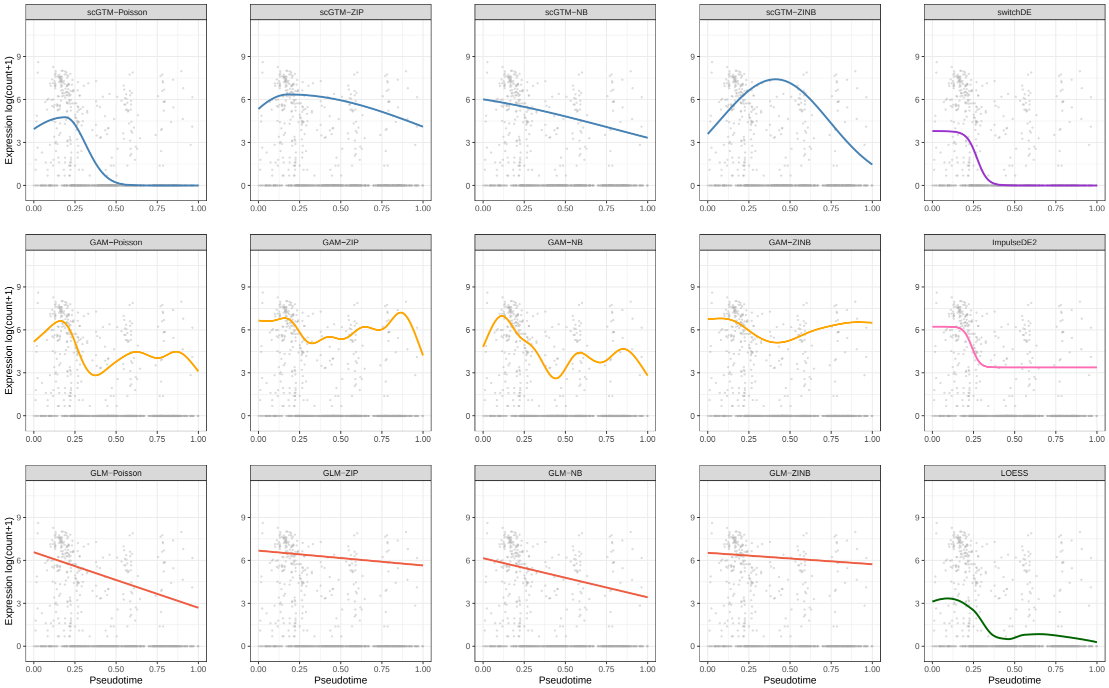


Figure S1: *PLAU*

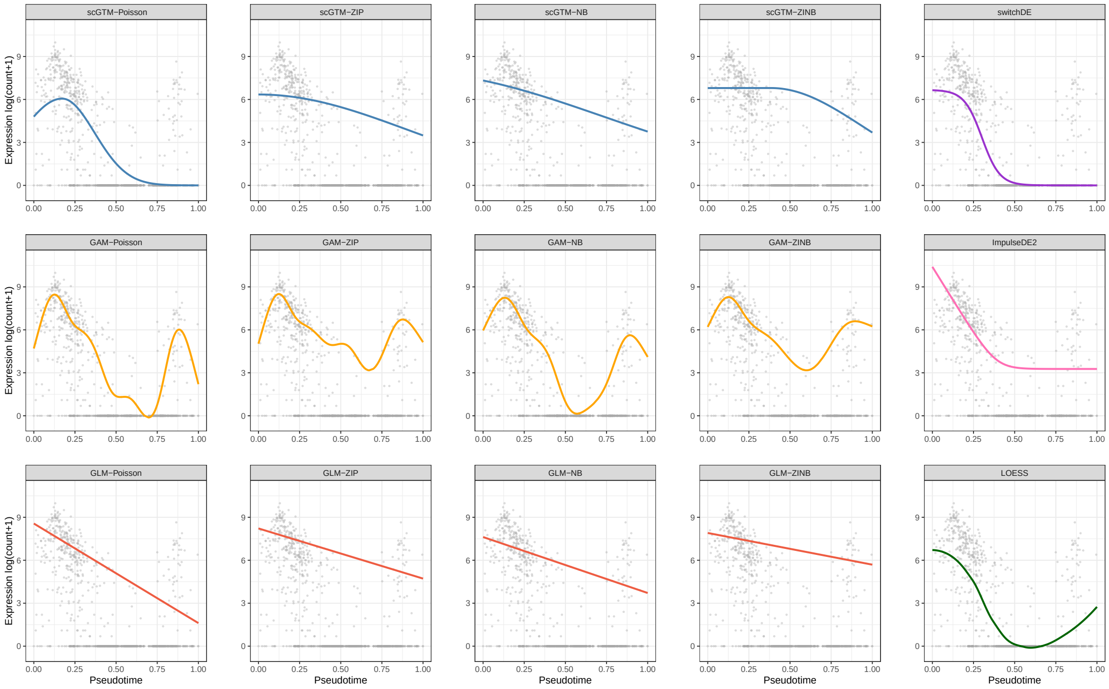


Figure S2: *MMP7*

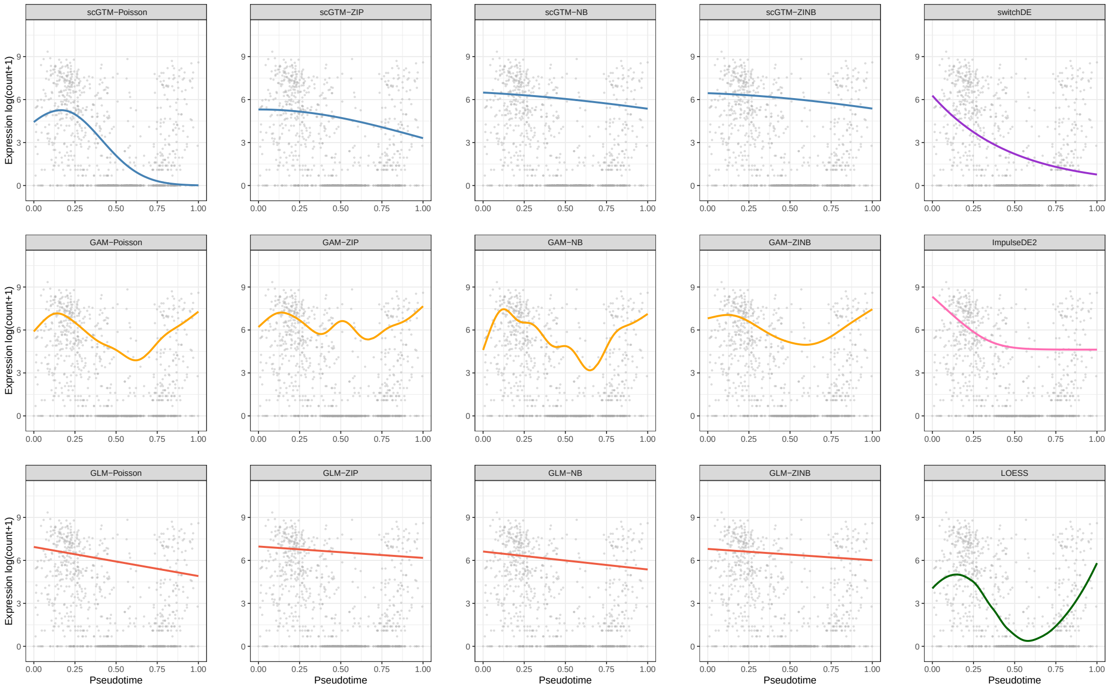


Figure S3: *THBS1*

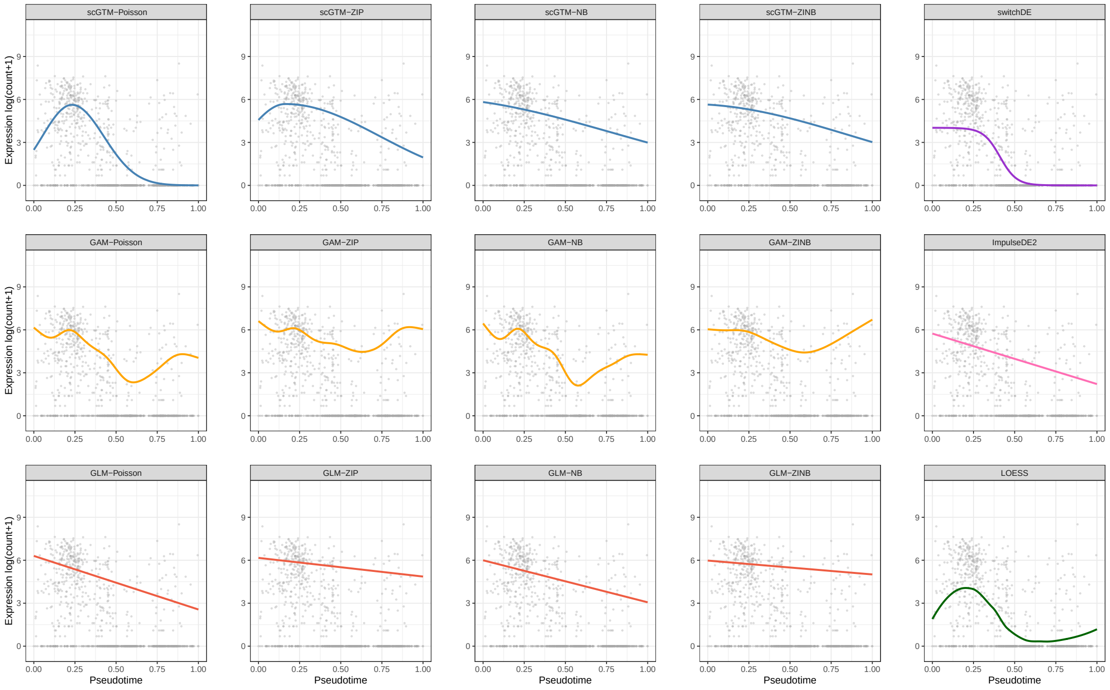


Figure S4: *CADM1*

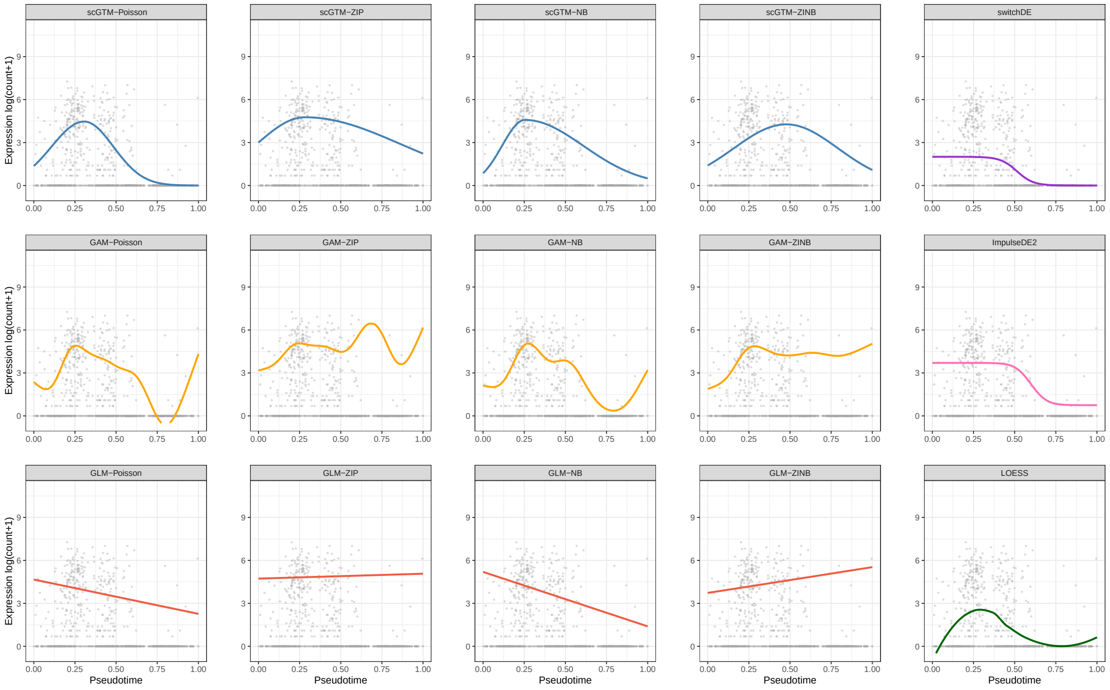


Figure S5: *NPAS3*

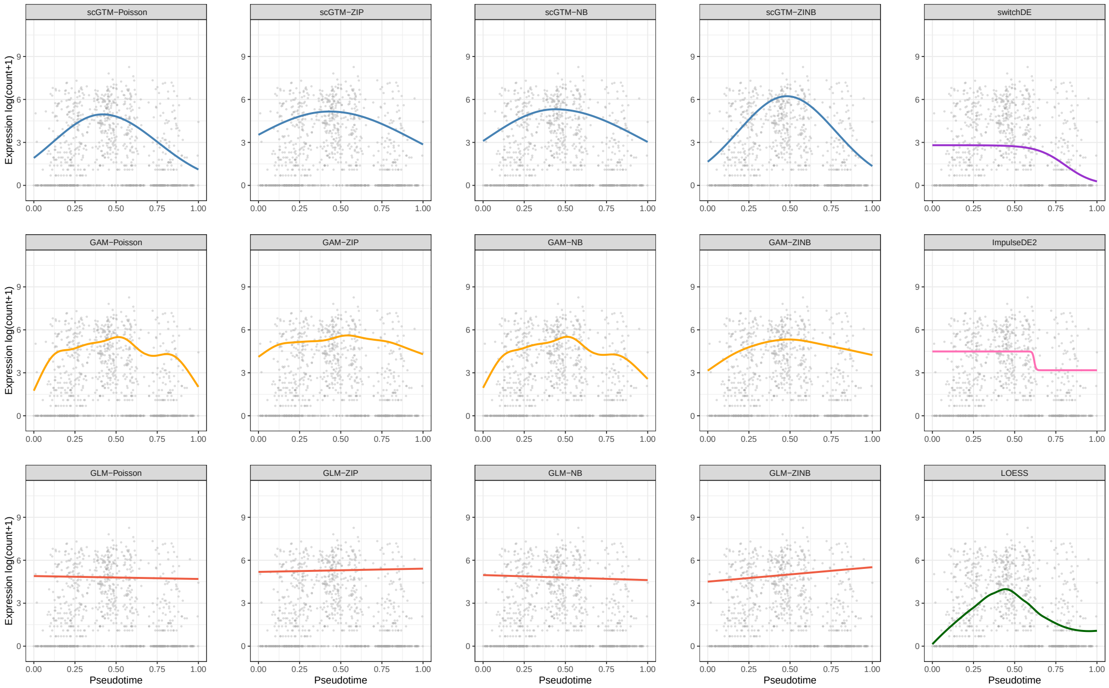


Figure S6: *ATP1A1*

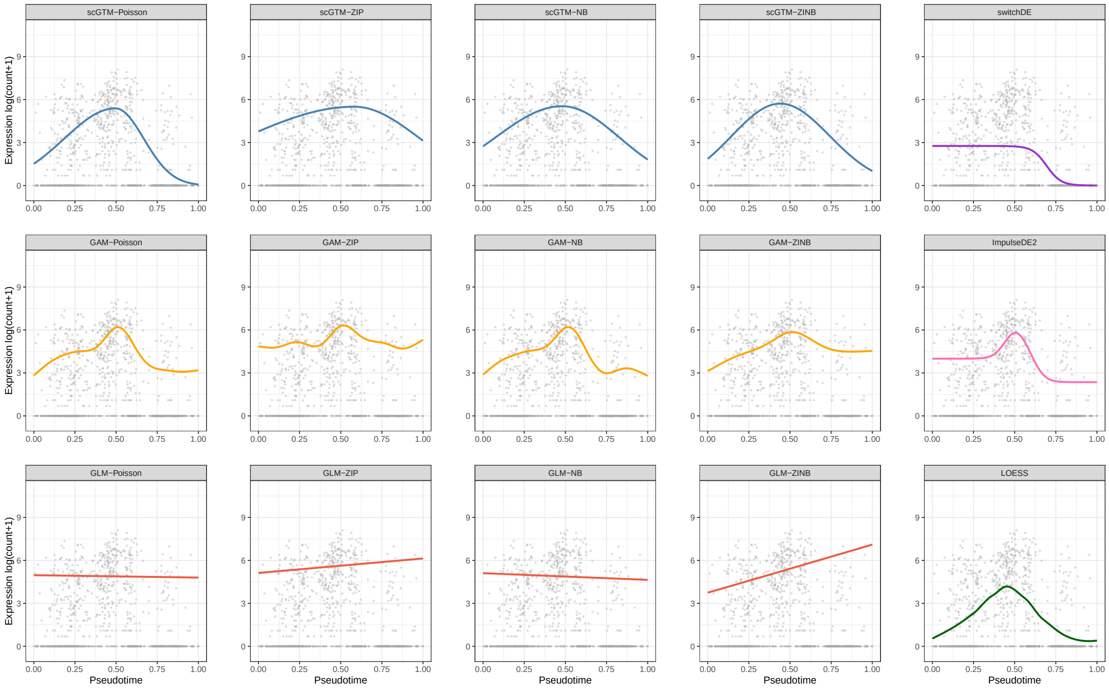


Figure S7: *ANK3*

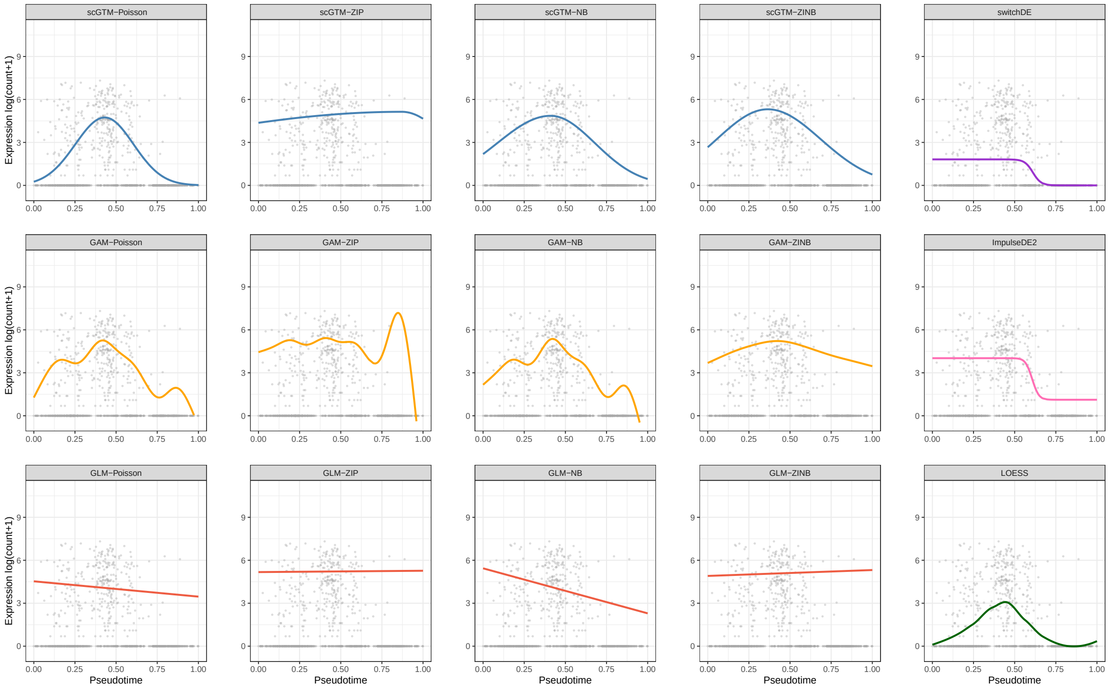


Figure S8: *ALPL*

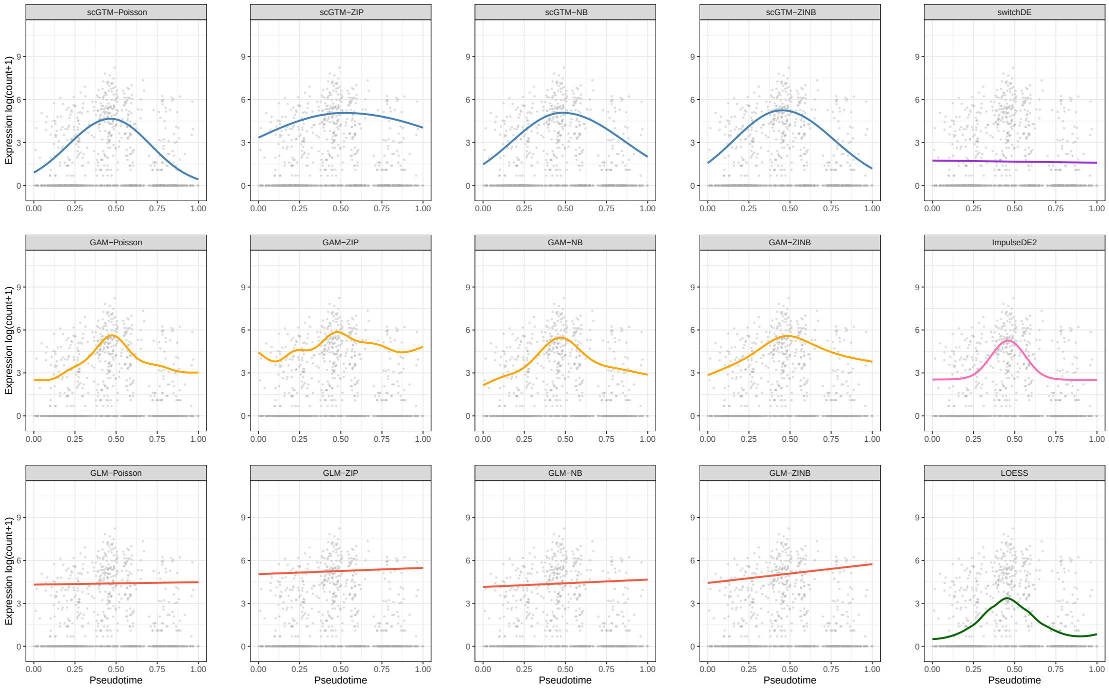


Figure S9: *TRAK1*

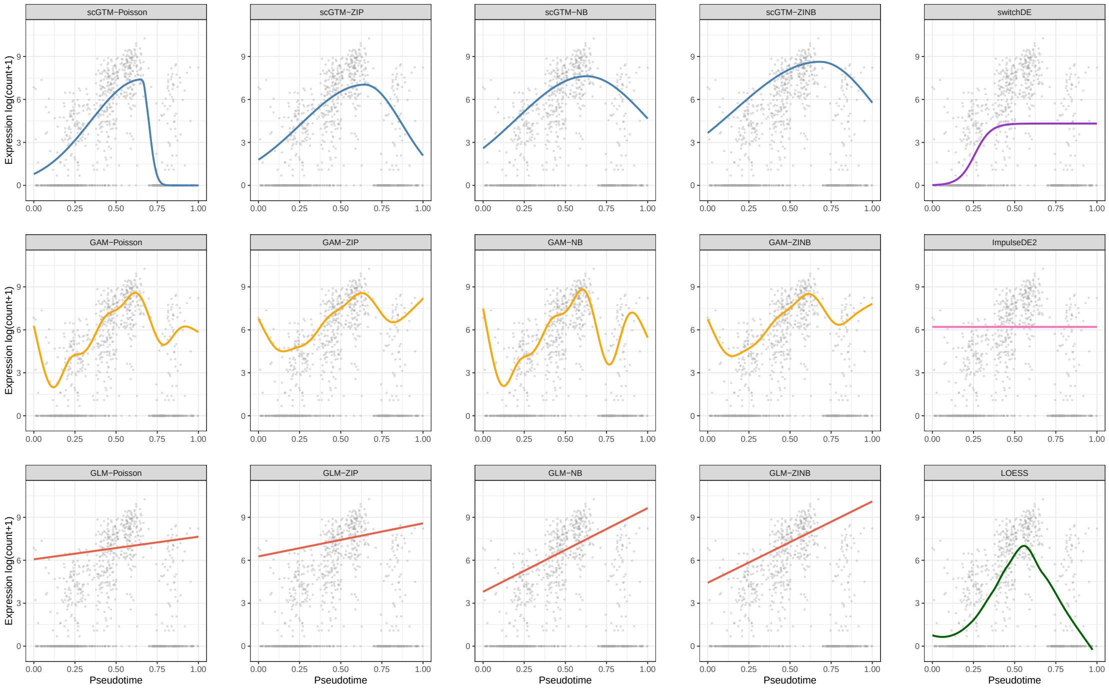


Figure S10: *SCGB1D2*

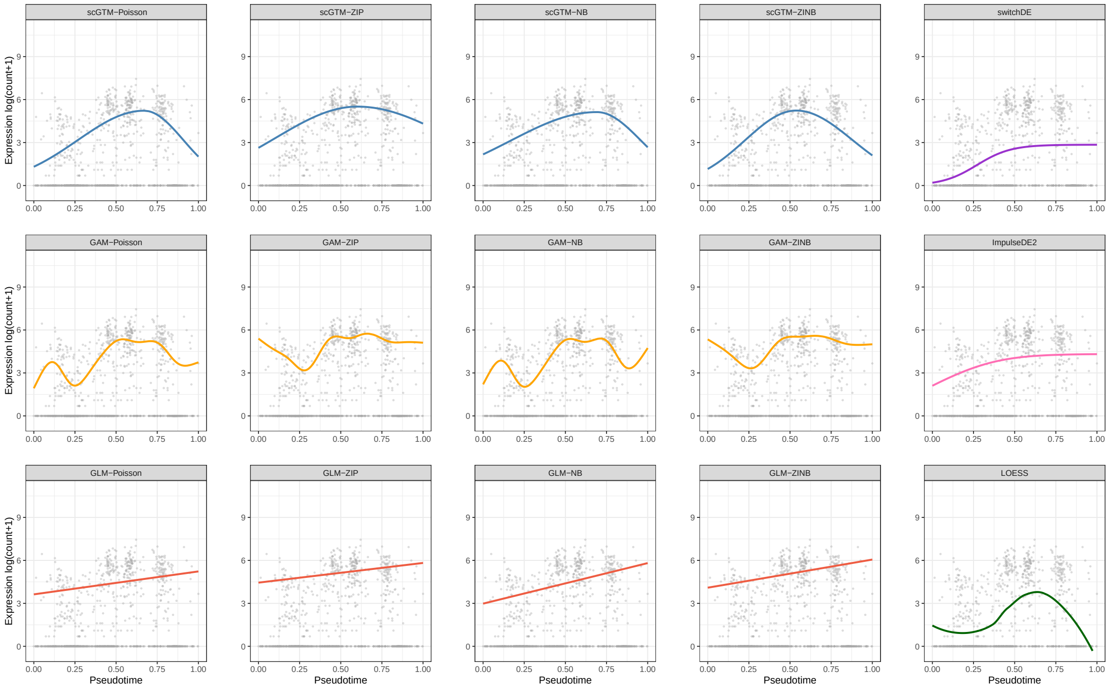


Figure S11: *MT1F*

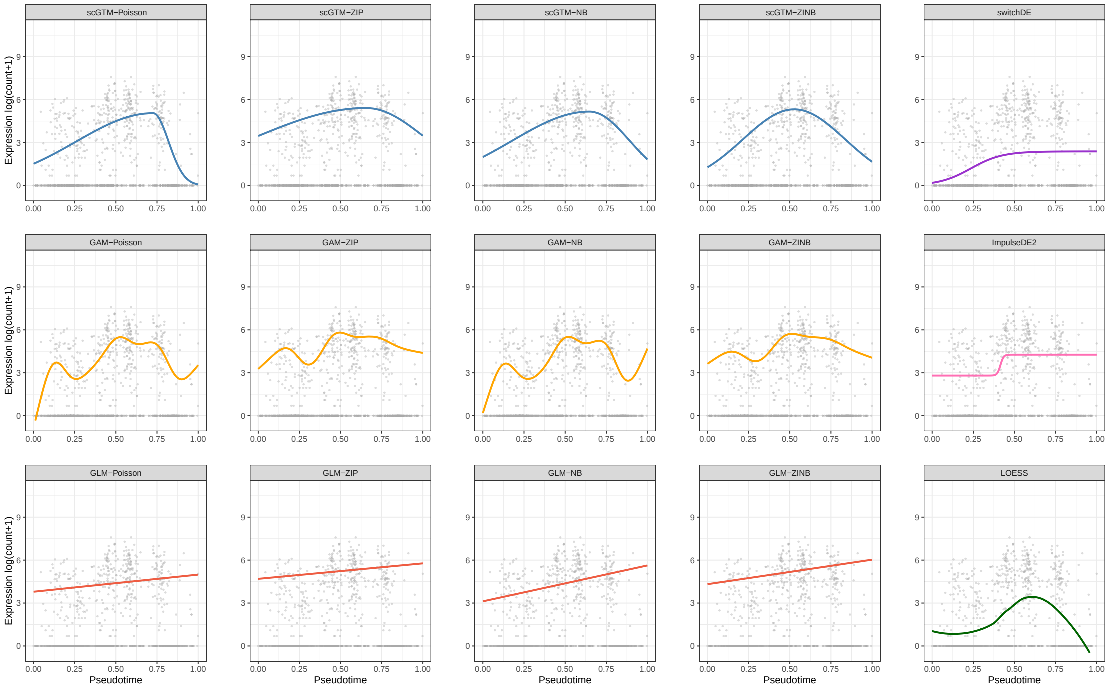


Figure S12: *MT1X*

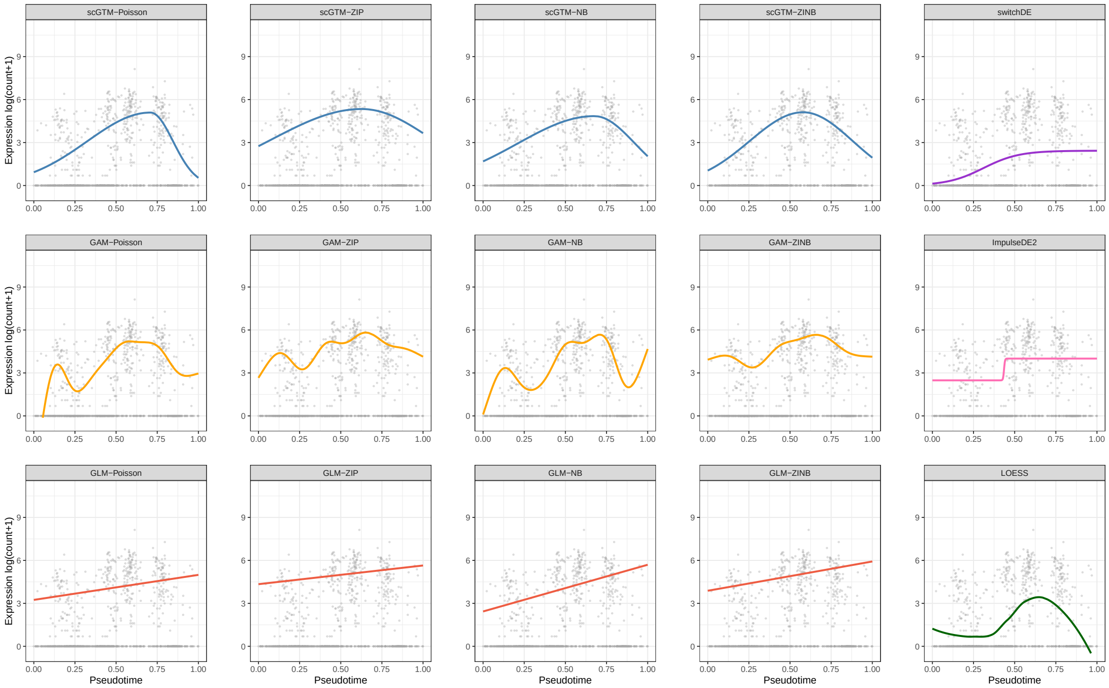


Figure S13: *MT1E*

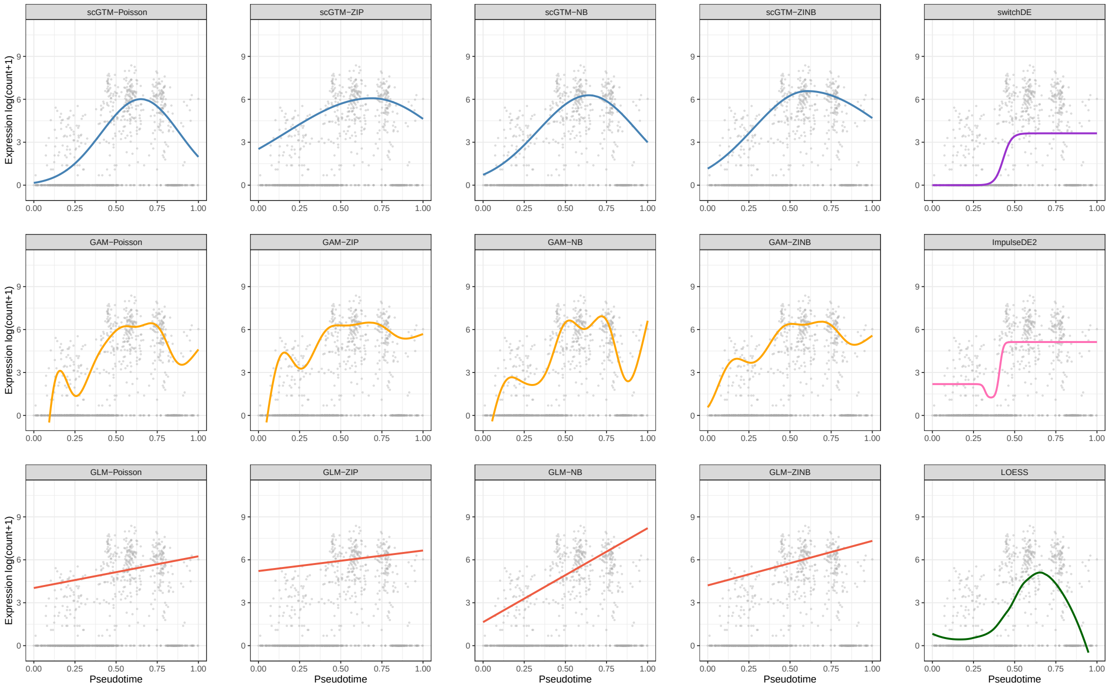


Figure S14: *MT1G*

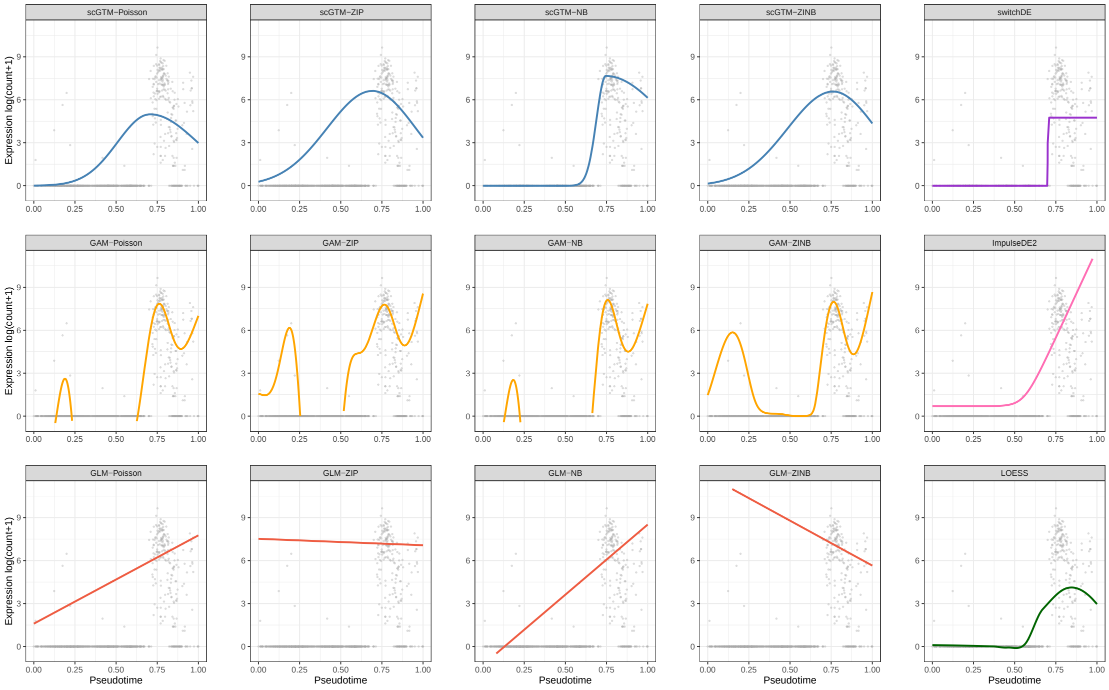


Figure S15: *CXCL14*

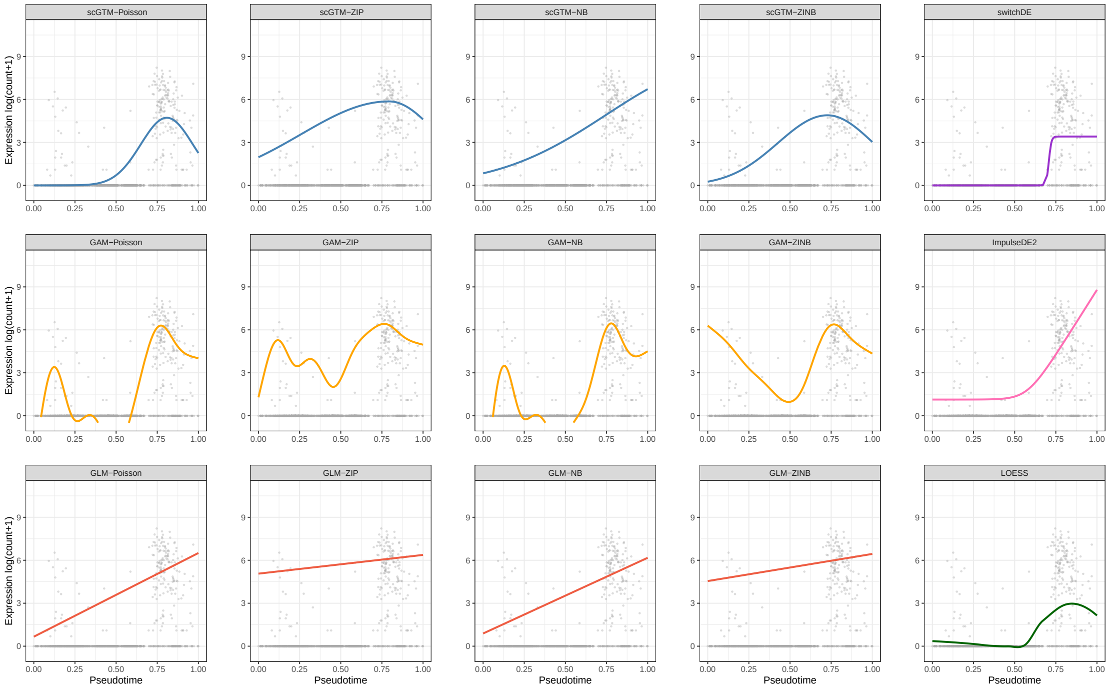


Figure S16: *DPP4*

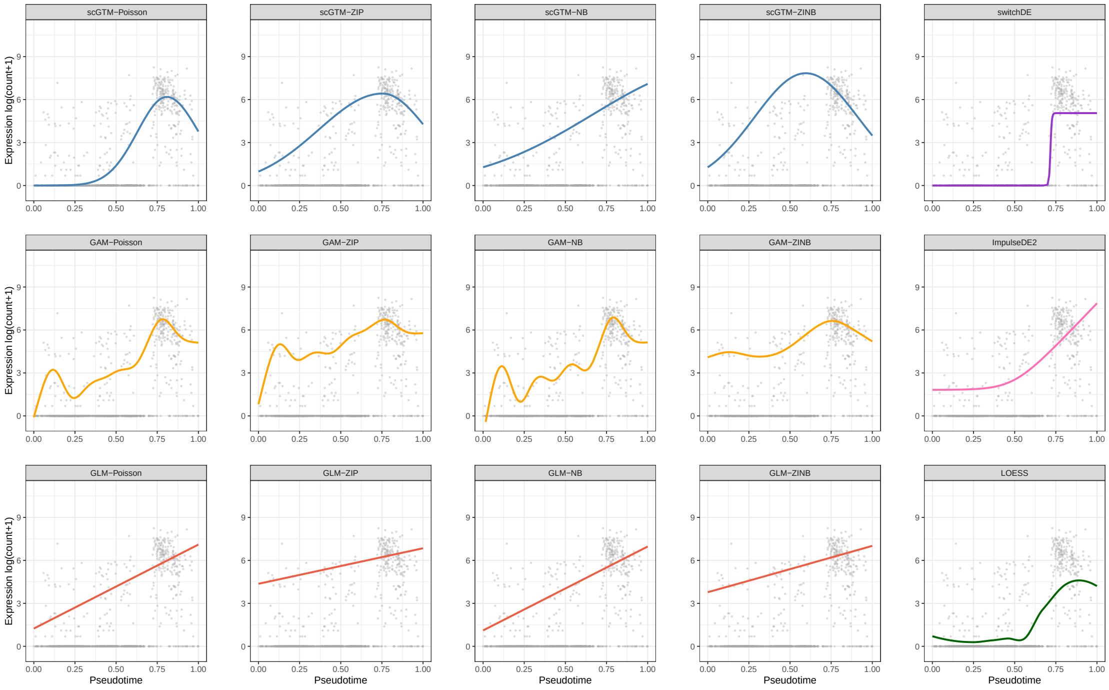


Figure S17: *NUPR1*

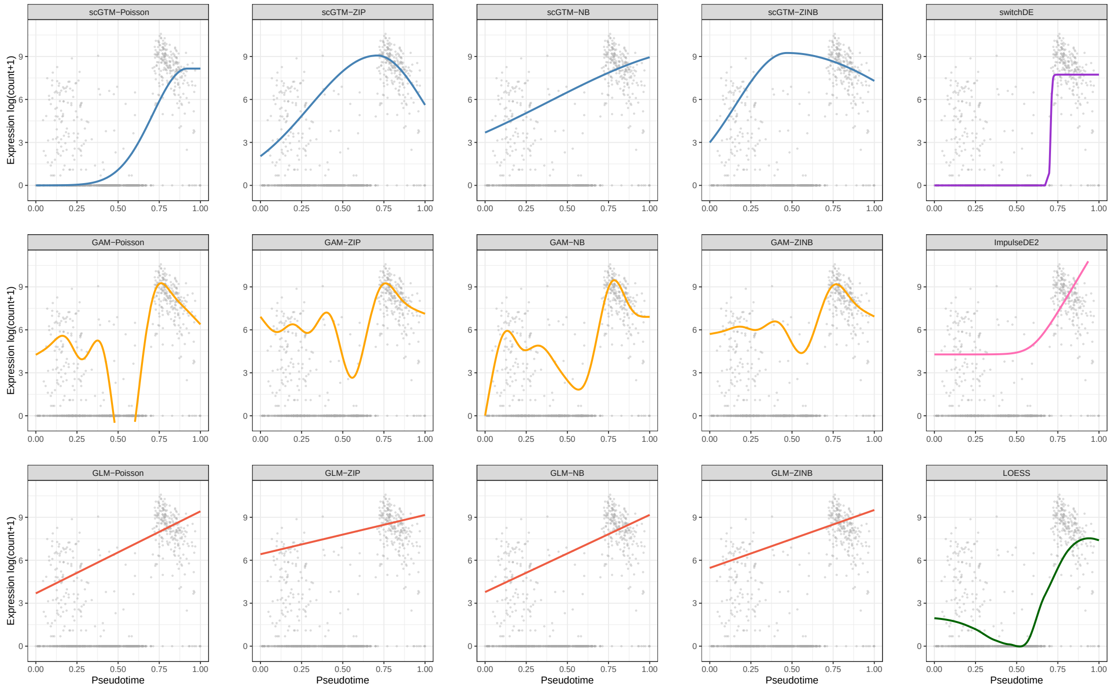


Figure S18: *GPX3*

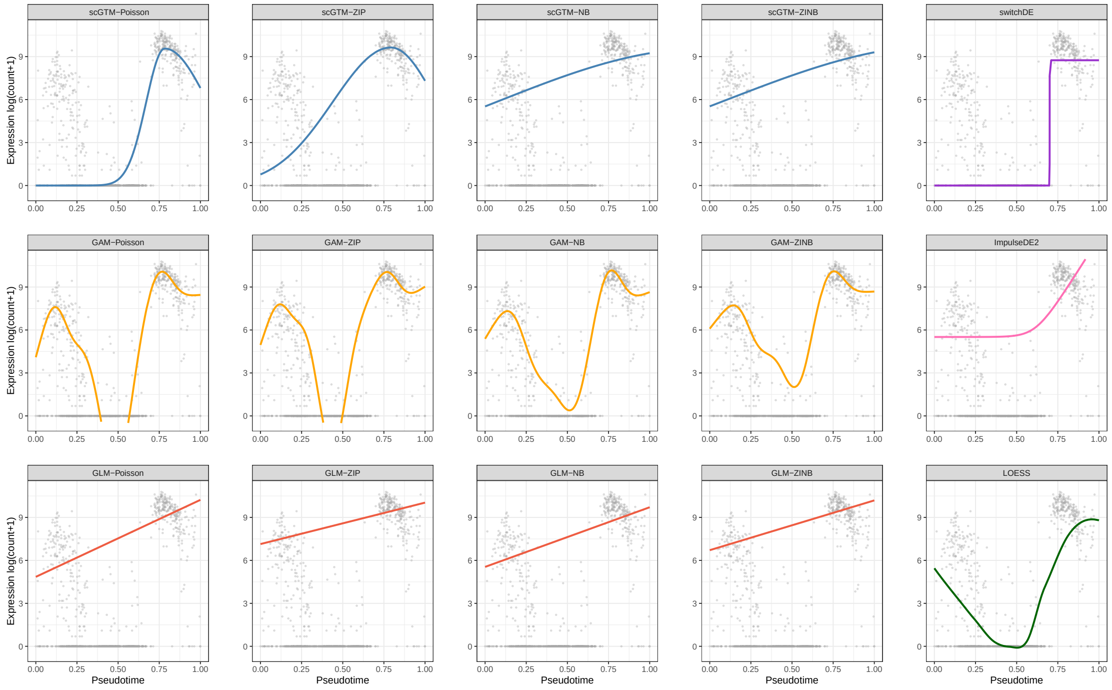


Figure S19: *PAEP*

#### S2 scGTM outperforms GAM and GLM in balancing goodness-of-fit and model complexity

We compare scGTM with GLM and GAM in terms of relative AIC. The relative AIC is calculated as follows: we first compute the AIC of all models for a particular gene and then divide all AIC values by the minimum AIC value so that the minimum value becomes 1; we call the resulting values the relative AIC values and write $\text{AIC}_{\text{rel}}^i = \text{AIC}^i / \min_j \text{AIC}^j$, where $i$ is the model index. In Fig. S20, we plot the boxplots of relative AIC values of different models on the 20 exemplar genes in the WANG dataset [Wang et al. (2020)]. From left to right, the four panels correspond to Poisson, ZIP, NB and ZINB respectively. Except for ZINB, scGTM has the top performance with other three distributions in terms of relative AIC. By the definition of AIC and relative AIC, the result suggests that scGTM is relatively robust to the choice of count distribution and does not have an overfitting problem.

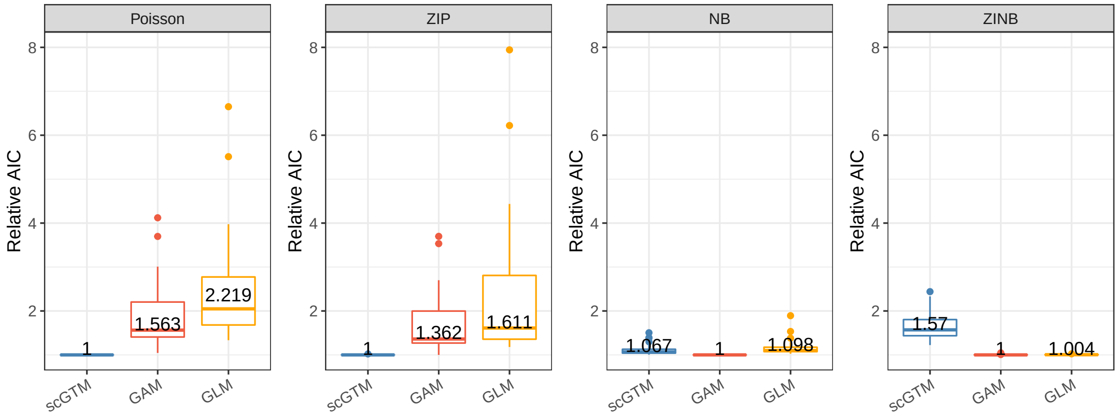


Figure S20: AIC comparison (balance of goodness-of-fit and model complexity) of scGTM with GAM and GLM on the WANG dataset [Wang et al. (2020)] (Supplementary Table S1). From left to right, the four panels correspond to the three models with the count distribution as Poisson, ZIP, NB, and ZINB, respectively. In each panel, from left to right, scGTM, GAM, and GLM are shown as blue, red, and orange boxplots, respectively; each boxplot shows the distribution of a model's relative AIC values across genes. A lower relative AIC value indicates better balance of goodness-of-fit and model complexity. With Poisson, ZIP, and NB as the count distribution (the left three panels), scGTM outperforms GAM and GLM.

### S3 Benchmarking scGTM against GAM by simulation and bootstrapping

We design a comprehensive simulation study to compare the scGTM with GAM. The simulation settings are summarized in Table S1. The design can be summarized in two aspects. The first aspect is the function of $\log(\tau_c + 1)$: the scGTM function, the quadratic function, and the logistic function; the latter two functions are used to show the robustness of the scGTM to trends that follow other functions. The second aspect is the trend shape: hill, valley, increasing, and decreasing. Together, we have eight function + shape settings, and we generate ten genes under each setting. Given the function of $\log(\tau_c + 1)$, we generate the gene expression count $Y_c$ from a negative binomial distribution with mean $\tau_c$.

Table S1: Overview of eight simulation settings.

| Function | Shape | Formula | Key parameter range(s) |
|---|---|---|---|
| scGTM | hill | $f(t_c) = \begin{cases} \mu_{\text{mag}} \exp\left(-k_1(t_c - t_0)^2\right) \text{ if } t_c \leq t_0 \\ \mu_{\text{mag}} \exp\left(-k_2(t_c - t_0)^2\right) \text{ if } t_c > t_0 \end{cases}$ | $0.4 < t_0 < 0.6$ |
| scGTM | valley | $f(t_c) = b - \begin{cases} \mu_{\text{mag}} \exp\left(-k_1(t_c - t_0)^2\right) \text{ if } t_c \leq t_0 \\ \mu_{\text{mag}} \exp\left(-k_2(t_c - t_0)^2\right) \text{ if } t_c > t_0 \end{cases}$ | $0.4 < t_0 < 0.6$ |
| scGTM | increasing | $f(t_c) = \begin{cases} \mu_{\text{mag}} \exp\left(-k_1(t_c - t_0)^2\right) \text{ if } t_c \leq t_0 \\ \mu_{\text{mag}} \exp\left(-k_2(t_c - t_0)^2\right) \text{ if } t_c > t_0 \end{cases}$ | $1 < t_0 < 1.2$ |
| scGTM | decreasing | $f(t_c) = \begin{cases} \mu_{\text{mag}} \exp\left(-k_1(t_c - t_0)^2\right) \text{ if } t_c \leq t_0 \\ \mu_{\text{mag}} \exp\left(-k_2(t_c - t_0)^2\right) \text{ if } t_c > t_0 \end{cases}$ | $-0.2 < t_0 < 0$ |
| quadratic | hill | $f(t_c) = b - \mu(t_c - t_0)^2$ | $0.4 < t_0 < 0.6$; $\mu > 0$ |
| quadratic | valley | $f(t_c) = b + \mu(t_c - t_0)^2$ | $0.4 < t_0 < 0.6$; $\mu > 0$ |
| logistic | increasing | $f(t_c) = \frac{\mu}{1+\exp(-k(t_c - t_0))}$ | $0.4 < t_0 < 0.6$; $\mu > 0$ |
| logistic | decreasing | $f(t_c) = \frac{-\mu}{1+\exp(-k(t_c - t_0))}$ | $0.4 < t_0 < 0.6$; $\mu > 0$ |

We first check if the scGTM can correctly recapitulate trend shapes. After fitting both hill- and valley-shaped scGTMs to a gene, we choose the model that has the smaller AIC value. Given the chosen model, we decide if a trend is monotone by checking if the confidence interval of $k_1$ or $k_2$ (Section 2.3) contains 0. For example, if a hill-shaped scGTM is chosen and the confidence interval of $k_1$ contains 0, we consider the trend as monotone decreasing; if a valley-shaped scGTM is chosen and the confidence interval of $k_1$ contains 0, we consider the trend as monotone increasing. Based on this decision process, the fitted scGTMs can perfectly distinguish between hill- and valley-shaped trends, and they have 97.5% accuracy for distinguishing increasing and decreasing trends. Given that a half of the genes are not simulated from the scGTM assumptions, these results demonstrate the robustness of scGTM.

We next check if the goodness-of-fit of the scGTM is comparable to that of GAM, which is designed to have great flexibility. For every gene $g$, we calculate the root mean square error $e_g$ between the true trend $f$ and the fitted trend $\hat{f}$:

$$e_g = \sqrt{\frac{\sum_{c=1}^{C} \left[f(t_c) - \hat{f}(t_c)\right]^2}{C}}\,.$$

A smaller $e_g$ means a better fit. We calculate the average of $G$ simulated genes' $e_g$'s and denote it as $\bar{e} = \frac{1}{G}\sum_{g=1}^{G} e_g$. For all 80 simulated genes (10 genes per setting × 8 settings), the scGTM performs similarly to GAM ($\bar{e}_{\text{scGTM}} = 0.080$; $\bar{e}_{\text{GAM}} = 0.077$). For the 40 genes simulated from the scGTM assumptions, the scGTM expectedly fits better than GAM does ($\bar{e}'_{\text{scGTM}} = 0.058$; $\bar{e}'_{\text{GAM}} = 0.075$). Fig. S21 shows one example gene per simulation setting, including the gene's true trend and the fitted trends by the scGTM and GAM. In particular, the scGTM fits increasing and decreasing trends better than GAM does.

Moreover, we use a bootstrap analysis to show that the fitted scGTM trend has a smaller variance than the fitted GAM trend does, at the cost of a larger bias. Fig. S22 shows the fitted scGTM and GAM trends

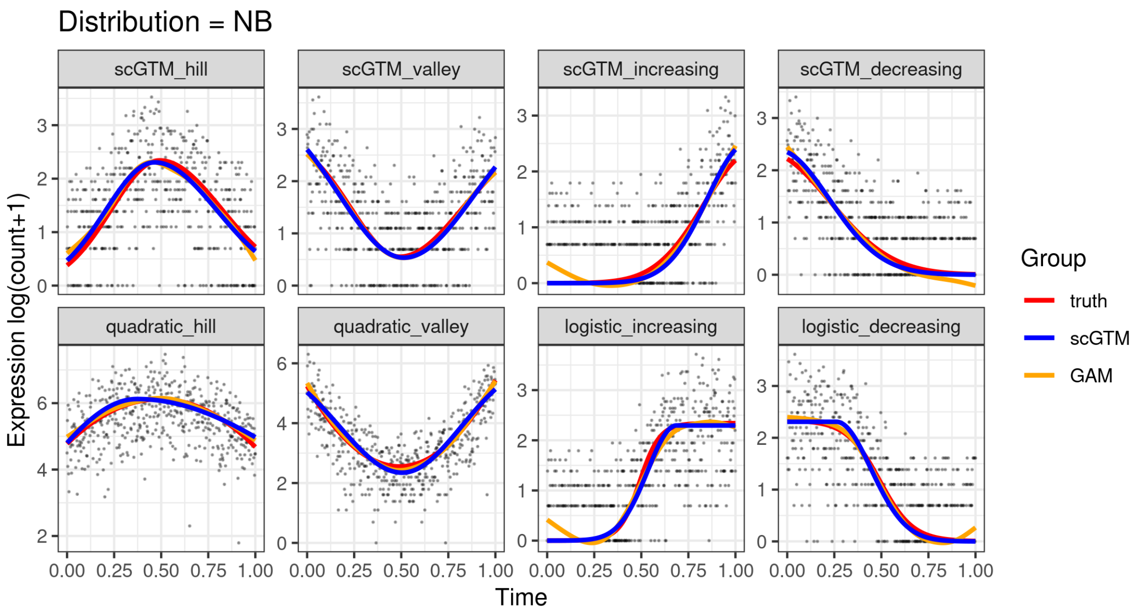


Figure S21: Comparison of scGTM with GAM for one example gene under each simulation setting.

on ten bootstrap samples of the *MAOA* gene in the WANG dataset [Wang et al. (2020)], and the scGTM trends are more stable across the bootstrap samples. This is more evident in Fig. S23, where the ten fitted trends for each model are overlaying with their mean trend; the root mean square error (between the fitted trends and the mean trend; calculated on 1000 evenly spaced pseudotime values in $[0, 1]$) is 0.399 for the scGTM and 2.799 for GAM.

Further, note that we already used the built-in penalization in the *mgcv* package to reduce the overfitting of GAM when we fit it (the *gam* function). Specifically, there is a smoothing parameter $\lambda$ to control the wiggliness of the fitted GAM trend, and $\lambda$ is estimated during the GAM fitting.

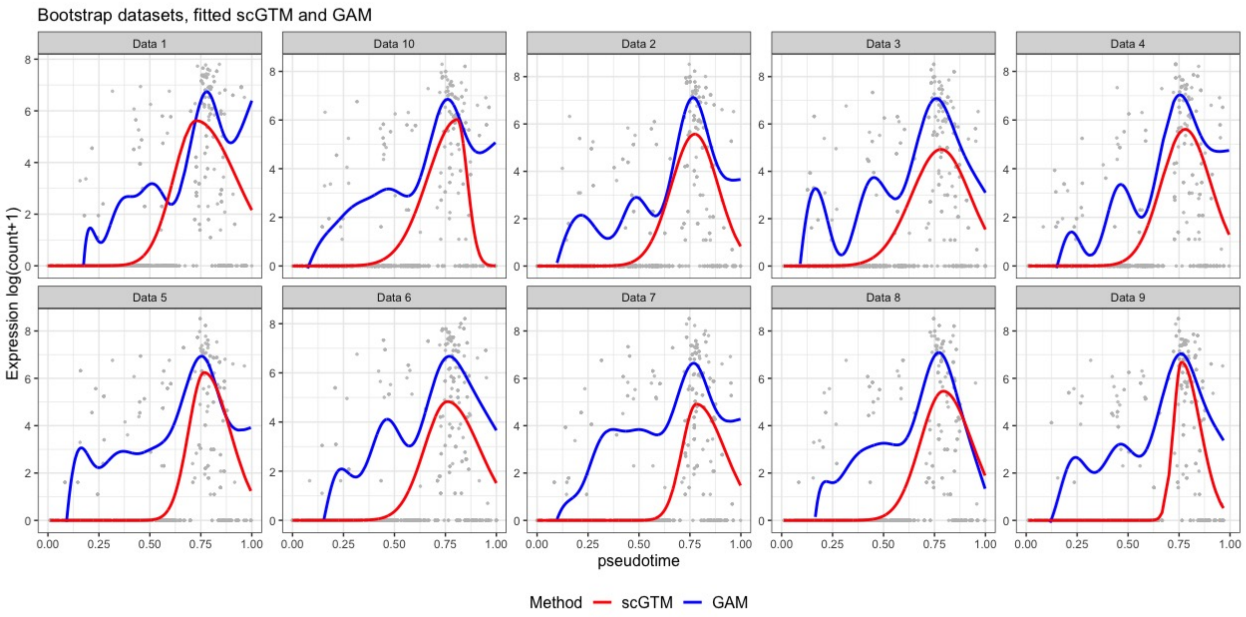


Figure S22: Fitted trends of scGTM and GAM on 10 bootstrap samples of the *MAOA* gene in the WANG dataset [Wang et al. (2020)].

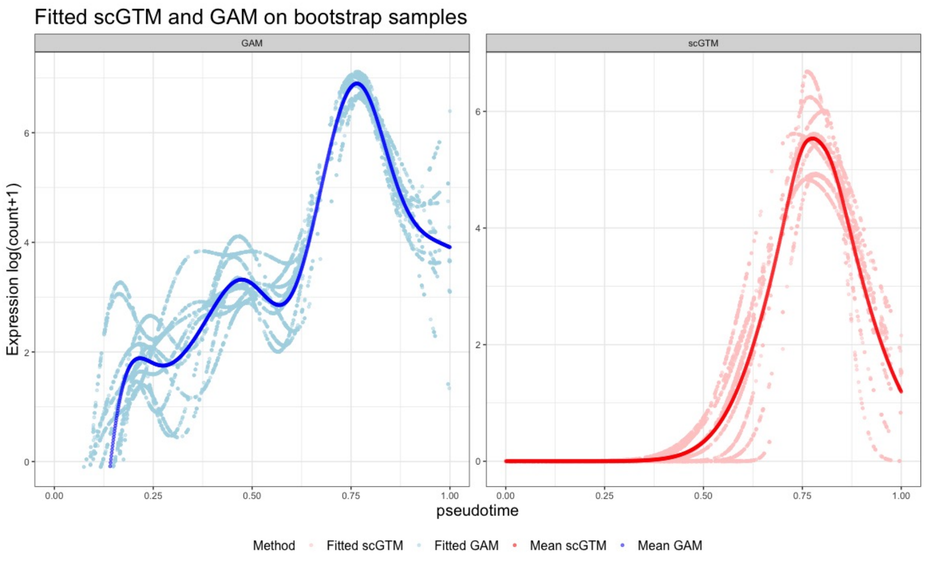


Figure S23: The fitted trends of scGTM and GAM on 10 bootstrap samples (light colored scatters) and the mean trends (dark colored curves) of the *MAOA* gene in the WANG dataset [Wang et al. (2020)].

### S4 scGTM is robust to pseudotime uncertainty

Unlike the observed (true) physical time, the pseudotime is inferred from data and thus intrinsically uncertain. The effects of pseudotime uncertainty on hypothesis testing (i.e., if a gene's expression changes with time) has been discussed in the PseudotimeDE method [Song and Li. (2022)]. However, the focus of this work is to proposed the scGTM for interpreting a trend, instead of testing whether a trend is different from a horizontal line, i.e., the focus of PseudotimeDE. Although scGTM does not directly account for the pseudotime uncertainty, we use simulation to show the robustness of scGTM to pseudotime uncertainty in a simple setting.

We first generate the true time $t_c$ and the true trend $f(t_c)$. To introduce uncertainty to $t_c$, we add normal random noise to obtain the pseudotime $t'_c = t_c + e$, where $e \sim N(0, 0.1^2)$. Fig. S24 shows that the scGTM fitted trends are still close to the true trend and correctly capture the trend shape.

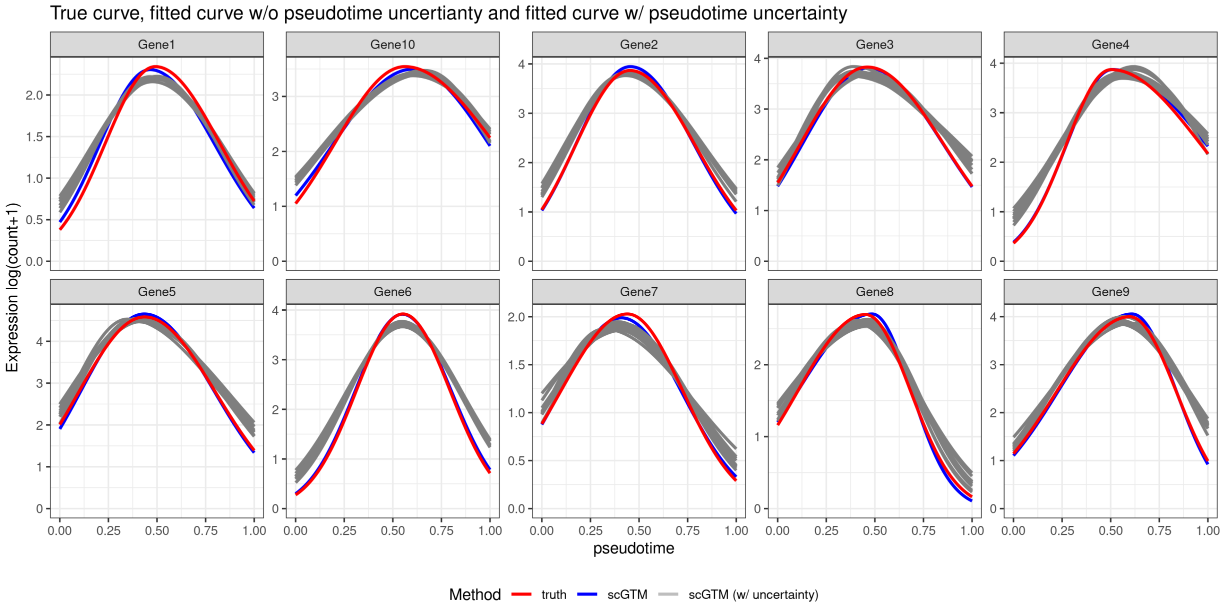


Figure S24: The effect of pseudotime uncertainty on scGTM fitting.

### S5 Derivation of Fisher information for confidence interval construction

Below we derive the Fisher information for the key parameters

$$\Theta^* = (\mu_{\text{mag}}, k_1, k_2, t_0)^\mathsf{T}$$

of the scGTM with Poisson distribution (for demonstration purposes; used for the results in Section 3.3). Recall the scGTM for a hill-shaped gene is

$$\begin{aligned} Y_c &\sim \text{Poisson}(\tau_c)\,, \\ \log(\tau_c + 1) &= \begin{cases} \mu_{\text{mag}} \exp\left(-k_1(t_c - t_0)^2\right) & \text{if } t_c \leq t_0 \\ \mu_{\text{mag}} \exp\left(-k_2(t_c - t_0)^2\right) & \text{if } t_c > t_0 \end{cases}. \end{aligned} \tag{S1}$$

The Fisher information for $\tau_c$ alone is $\mathcal{I}_{\text{Poi}}(\tau_c) = 1/\tau_c$, $c = 1, \ldots, C$, and every $\tau_c$ is related to $\Theta^*$ via (S1). Then by the chain rule, the Fisher information for $\Theta^*$ is

$$\begin{aligned} \mathcal{I}_{\text{Poi}}(\Theta^*) &= \sum_{\{c:\ t_c \leq t_0\}} \left(1 + \frac{1}{\exp(f_1) - 1}\right) \boldsymbol{x}_c \boldsymbol{x}_c^\mathsf{T} + \sum_{\{c:\ t_c > t_0\}} \left(1 + \frac{1}{\exp(f_2) - 1}\right) \boldsymbol{x}_c \boldsymbol{x}_c^\mathsf{T}\,, \\ \text{where} \quad \boldsymbol{x}_c &= \begin{cases} (f_1/\mu_{\text{mag}}, \quad (t_c - t_0)^2 f_1, \quad 0, \quad 2k_1(t_c - t_0) f_1)^\mathsf{T} & \text{if } t_c \leq t_0 \\ (f_2/\mu_{\text{mag}}, \quad 0, \quad (t_c - t_0)^2 f_2, \quad 2k_2(t_c - t_0) f_2)^\mathsf{T} & \text{if } t_c > t_0 \end{cases}, \\ f_1 &= \mu_{\text{mag}} \exp\left(-k_1(t_c - t_0)^2\right), \\ f_2 &= \mu_{\text{mag}} \exp\left(-k_2(t_c - t_0)^2\right). \end{aligned} \tag{S2}$$

Then the estimated asymptotic covariance of $\widehat{\Theta}^*$ is $\widehat{\mathcal{I}}_{\text{Poi}}^{-1}(\widehat{\Theta}^*)$.

### S6 Particle swarm optimization

Swarm intelligence algorithms, such as ant colony algorithms [Dorigo et al. (2006)], cuckoo search algorithms, [Yang and Deb (2009)] and firefly algorithms [Yang (2009)], mimic the behaviour of a swarm to solve optimization problems. They are now receiving more and more interest and attention not only in the literature of mathematics, but also in econometrics, optimal design, engineering, etc [Yang(2017)]. Particle swarm optimization (PSO), proposed by Kennedy and Eberhart in 1995 [Kennedy and Eberhart (1995)], is one of the most widely used swarm intelligence algorithms to optimize an objective function with boundary constraints. It is the main optimization tool in this paper and is introduced in below.

PSO solves an optimization problem by producing a sequence of candidate solutions. Unlike the gradient descent algorithms widely used for deep learning, PSO does not require either differentiability or convexity [Boyd et al. (2004)] of the objective function and constraints. Therefore, PSO is particularly useful when the objective function does not have desirable analytical properties (e.g., not differentiable).

PSO encodes swarm intelligence, such as bird flocking, into two simple dynamic equations to solve optimization problems of the following form:

$$\min f(\boldsymbol{x}) \quad \text{s.t. } \boldsymbol{x} \in \mathcal{S}\,,$$

where $\boldsymbol{x} \in \mathbb{R}^d$ is a $d$-dimensional vector, $f(\boldsymbol{x})$ is a real-valued objective function (measurability is the only requirement), and $\mathcal{S} \subset \mathbb{R}^d$ is the search space or domain of $\boldsymbol{x}$). The algorithm starts with $n$ candidate values of $\boldsymbol{x}$, denoted as $\boldsymbol{x}_1^0, \cdots, \boldsymbol{x}_n^0$. Each $\boldsymbol{x}_i^0$, $i = 1, \cdots n$, represents a *particle* and is initialized with a velocity vector $\boldsymbol{v}_i^0 \in \mathbb{R}^d$. Then for $i = 1, \cdots, n$, PSO iterates with the following two equations [Bratton and Kennedy(2007)]:

$$\begin{aligned} \boldsymbol{v}_i^{k+1} &= w\boldsymbol{v}_i^k + c_1 r_{i1}^k(\widehat{\boldsymbol{x}}_i^k - \boldsymbol{x}_i^k) + c_2 r_{i2}^k(\widehat{\boldsymbol{x}}^k - \boldsymbol{x}_i^k)\,, \\ \boldsymbol{x}_i^{k+1} &= \boldsymbol{x}_i^k + \boldsymbol{v}_i^{k+1}\,, \end{aligned} \tag{S3}$$

where $k = 0, 1, \cdots$ is the number of iterations finished, $w$ is called the *inertia weight*, $c_1$ and $c_2$ are called the *cognitive* and *social* parameters respectively, and $r_{i1}^k$ and $r_{i2}^k$ are two random numbers independently generated uniformly from $[0, 1]$. Usually, $w, c_1$, and $c_2$ are set to numbers in $[0, 2]$ by users. Most importantly,

$$\begin{aligned} \widehat{\boldsymbol{x}}_i^k &= \arg\min_{\boldsymbol{x} \in \mathcal{A}_i} f(\boldsymbol{x})\,, \\ \widehat{\boldsymbol{x}}^k &= \arg\min_{\boldsymbol{x} \in \cup_{i=1}^n \mathcal{A}_i} f(\boldsymbol{x})\,, \\ \text{where } \mathcal{A}_i &= \{\boldsymbol{x}_i^t : t = 0, \cdots, k\}\,. \end{aligned}$$

Thus, $\widehat{\boldsymbol{x}}_i^k$ is the best position recorded by particle $i$ up to the $k^{\text{th}}$ iteration, and $\widehat{\boldsymbol{x}}^k$ is the best position recorded by the whole swarm up to the $k^{\text{th}}$ iteration. The inertia weight $w$ controls the level of a particle moving towards its last direction $\boldsymbol{v}_i^k$. The cognitive parameter $c_1$ represents how a particle is affected by its best known position $\widehat{\boldsymbol{x}}_i^k$. Similarly, the social parameter $c_2$ determines the influence of the swarm's best knowledge $\widehat{\boldsymbol{x}}^k$ on particle $i$. Because $\widehat{\boldsymbol{x}}^k$ is the best solution found by the whole swarm, the set of equations (S3) is also called the *global best PSO* [Bratton and Kennedy(2007)].

To better understand the logic of PSO, suppose we have 10 ants starting around origin $(0, 0)$ and they are looking for food at point $(2, 2)$ (left panel of Fig. S25). Background colors represent distances to the food. The initial position of each ant corresponds to $\boldsymbol{x}_i^0$, and the objective function $f(\boldsymbol{x})$ is the Euclidean distance between point $\boldsymbol{x}$ and the food at $(2, 2)$. Each ant is initialized with an velocity vector $\boldsymbol{v}_i^0$ (blue arrow). After moving one step, ants re-analyze their positions and distances to the food so that (1) the best position of ant $i$ is recorded as $\widehat{\boldsymbol{x}}_i^1$; (2) the best position of all ants is recorded as $\widehat{\boldsymbol{x}}^1$. Here the *best* position has the minimum distance to the food at (2,2). Then, each ant re-corrects its velocity according to equation (S3) (middle panel of Fig. S25). After several iterations, all ants gather around the food and the velocity decreases to 0 gradually (right panel of Fig. S25).

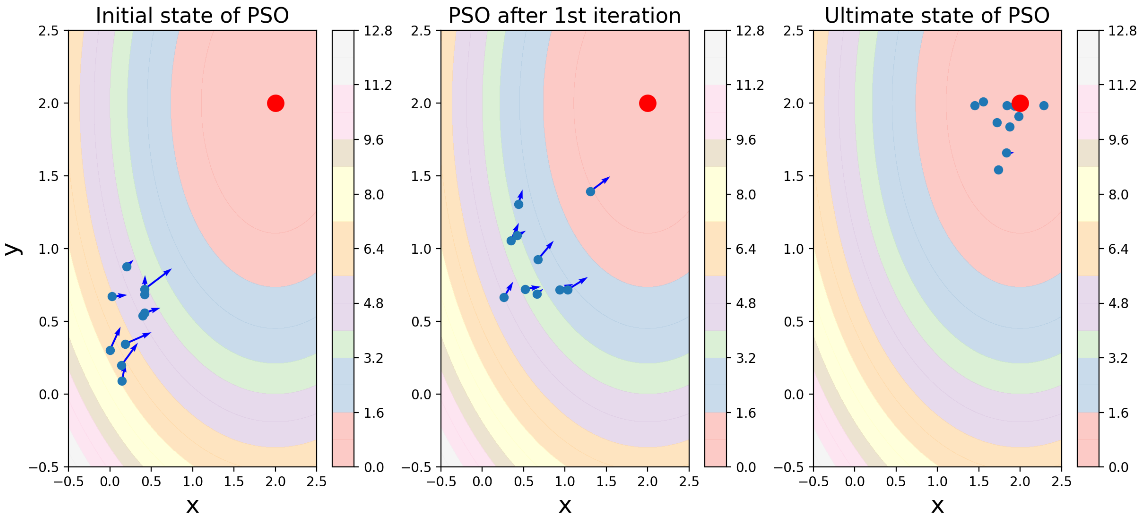


Figure S25: Illustration of PSO.

### S7 Datasets, R packages, and R functions used in this paper

Table S2: Overview of datasets used.

| Dataset | Sequencing protocol | Gene # | Cell # | Description | Ref |
|---|---|---|---|---|---|
| LPS | Fluidigm c1 | 4018 | 390 | mouse bone-marrow-derived dendritic cells after stimulation with LPS | Shalek et al. (2014) |
| WANG | Fluidigm C1 | 22036 | 984 | human unciliated epithelia cells during the menstrual cycle | Wang et al. (2020) |
| GYRUS | 10x Genomics Chromium | 2291 | 678 | mouse developing dentate gyrus | Hochgerneret al.(2018) |

Table S3: Overview of R packages and functions used for fitting GLMs and GAMs.

| Model | R package | R function[1] | Parameter `family`[2] |
|---|---|---|---|
| GLM-Poisson | `stats` | `glm()` | `poisson()` |
| GLM-ZIP | `mgcv` | `gam()` | `ziP()` |
| GLM-NB | `mgcv` | `gam()` | `negbin()` |
| GLM-ZINB | `zigam` | `zinbgam()` | |
| GAM-Poisson | `mgcv` | `gam()` | `poisson()` |
| GAM-ZIP | `mgcv` | `gam()` | `ziP()` |
| GAM-NB | `mgcv` | `gam()` | `negbin()` |
| GAM-ZINB | `zigam` | `zinbgam()` | |

[1] The R function to call in the R package.

[2] The `family` parameter (i.e., distribution) to specify in the R function. For example, `ziP()` refers to the ZIP distribution and can be specified in the `gam()` function in the `mgcv` package. There is no such parameter in the `zinbgam()` function in the `zigam` package.

### S8 Additional detail of analysis in the paper

#### S8.1 Pseudotime inference

For the LPS and GYRUS datasets, we use the R package `slingshot` (version 2.0.0) to infer cell pseudotime. We use the top 2 principal components on the $\log(\text{count} + 1)$ matrix as the input of `slingshot`. For the WANG dataset, the pseudotime is provided by the authors of the original study [Wang et al. (2020)].

#### S8.2 GO analysis

We use the R package `clusterProfiler` (4.0.5) to perform GO analysis in Section 3.3. We set the $p$-value cutoff and $q$-value cutoff as 0.01 and 0.05, respectively. We set the ontology type as "BP (Biological Process)". We use the function `clusterProfiler::simplify` to further reduce the redundancy in GO terms.

#### S8.3 Visualization

Most figures are made with the R package `ggplot2` (version 3.3.5). Figure 5 is generated by the R package `ComplexHeatmap` (version 2.9.3).

#### S9 Selection of trend shapes for the genes in Fig. 1b–c

Fig. S26 shows alternatively fitted trends (valley-shaped trend for *Tmsb10* and hill-shaped trend for *NFKBIA*) for the two genes in Fig. 1b–c. The AIC values show that the hill-shaped trend in Fig. 1b fits better for *Tmsb10*, and the valley-shaped trend in Fig. 1c fits better for *NFKBIA*.

| Gene name | Trend | AIC |
|---|---|---|
| *Tmsb10* | Hill-shaped | 16103.7 |
| *Tmsb10* | Valley-shaped | 78571.0 |
| *NFKBIA* | Hill-shaped | 161805.2 |
| *NFKBIA* | Valley-shaped | 13345.1 |

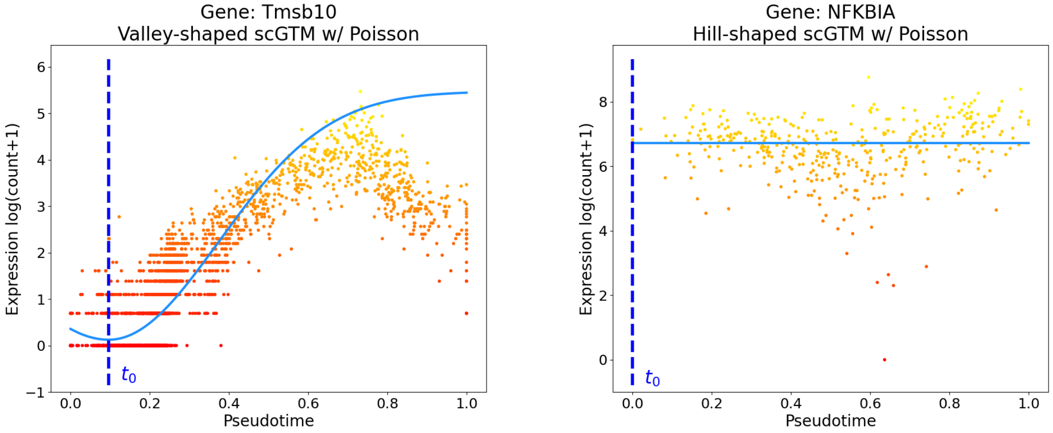


Figure S26: Alternatively fitted trends for the genes *Tmsb10* and *NFKBIA* in Fig. 1b–c.

#### S10 scGTM extension can capture more complicated gene trends

Currently the scGTM is designed for detecting simple and easy-to-interpret gene expression trends including monotone, hill-shaped and valley-shaped trends. The reason is that most genes of biological interests are observed to follow one of these simple trends. We deem this reasonable because each pseudotime trajectory is expected to indicate a directional change process, such as development and immune response, along which important genes usually have no more than one hill or valley.

Meanwhile, in practice some genes may exhibit more complicated patterns. Accordingly, the scGTM is extendable by assuming a more complicated mean function, whose estimation can still be achieved by the PSO algorithm (whose major advantage is its flexibility). To demonstrate this functionality of the scGTM, we conduct a simulation study where we use the sine function to generate one gene's true expression trend along the pseudotime. With its mean function set as as the sine function, the scGTM accurately estimates the gene trend (Fig. S27).

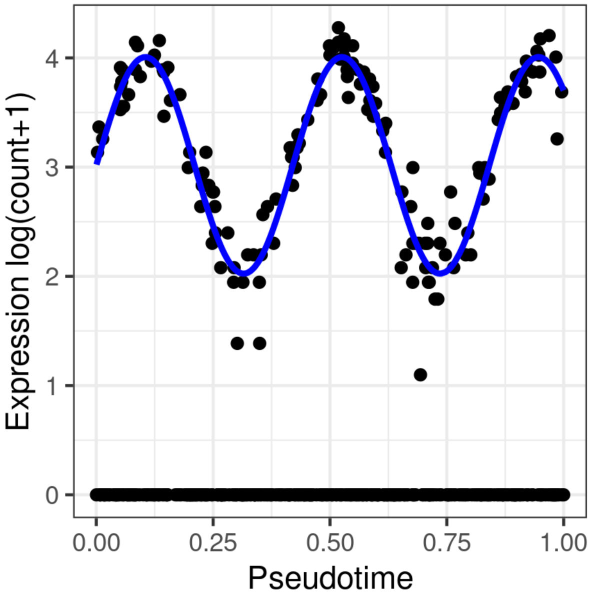


Figure S27: scGTM extension correctly captures the sine gene trend.

### S11 Applying scGTM on 1,382 human menstrual cycle genes

We choose the 20 genes because they were analyzed and provided with biological interpretations in [Wang et al. (2020)]. The complete dataset contains 22,036 genes, most of which are not related to human menstrual cycle and thus do not exhibit notable expression trends. To further show the performance of scGTM, we focused on the 1,382 menstrual cycle genes reported in [Wang et al. (2020)], and we applied the scGTM to fit these genes' expression trends. The results are still satisfying.

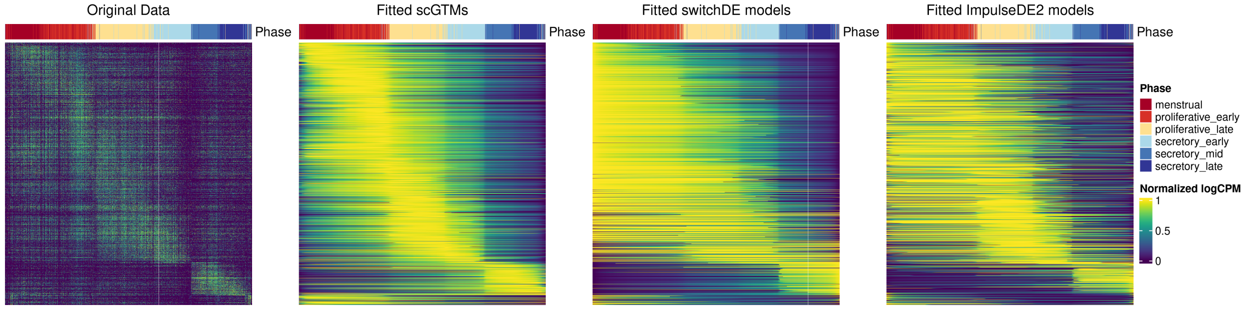


Figure S28: Fitted expression trends by scGTM, switchDE, and ImpulseDE2 for 1,382 menstrual cycle related genes in [Wang et al. (2020)].
